# Mitochondrial dysfunction drives a neuronal exhaustion phenotype in methylmalonic aciduria

**DOI:** 10.1101/2024.03.15.585183

**Authors:** Matthew C.S. Denley, Monique S. Straub, Giulio Marcionelli, Miriam A. Güra, David Penton, Igor Delvendahl, Martin Poms, Beata Vekeriotaite, Sarah Cherkoui, Federica Conte, Ferdinand von Meyenn, D. Sean Froese, Matthias R. Baumgartner

## Abstract

Methylmalonic aciduria (MMA) is an inborn error of metabolism resulting in loss of function of the enzyme methylmalonyl-CoA mutase (MMUT). Despite acute and persistent neurological symptoms, the pathogenesis of MMA in the central nervous system is poorly understood, which has contributed to a dearth of effective brain specific treatments. Here we utilised patient-derived induced pluripotent stem cells and *in vitro* differentiation to generate a human neuronal model of MMA. We reveal strong evidence of mitochondrial dysfunction caused by deficiency of MMUT in patient neurons. By employing patch-clamp electrophysiology, targeted metabolomics, and bulk transcriptomics, we expose an altered state of excitability, which is exacerbated by application of 2-dimethyloxoglutarate, and we suggest may be connected to metabolic rewiring. Our work provides first evidence of mitochondrial driven neuronal dysfunction in MMA, which through our comprehensive characterisation of this paradigmatic model, enables first steps to identifying effective therapies.

## Introduction

Mitochondrial diseases are a diverse group of genetic disorders that lead to dysfunctional mitochondria and energetic depletion. Pathogenic variants in approximately 1136 genes which encode mitochondrial-resident proteins^1^, found on both nuclear and mitochondrial DNA, may cause mitochondrial diseases. Together, 23 per 100,000 people harbour pathogenic variants that have or will convey a mitochondrial disease^2^. However, the inherent clinical and genetic heterogeneity of mitochondrial diseases make them difficult to diagnose, hard to treat, and complex to study^3,4^. Nevertheless, mitochondrial diseases affecting components of oxidative phosphorylation (OXPHOS) or proteins proximal to the tricarboxylic acid cycle (TCA cycle) often report overlapping symptoms^5,6^, particularly within the central nervous system (CNS)^7,8^.

Isolated methylmalonic aciduria (MMA) is an archetypical mitochondrial disease caused by pathogenic variants leading to the absence or deficiency of the nuclear-encoded mitochondrial matrix-residing enzyme methylmalonyl-CoA mutase (MMUT; MIM #251000)^9^. It is typified by accumulation of methylmalonic acid and related compounds in blood and urine; however, increased levels of these disease-related compounds are likely not sufficient to explain overall clinical presentation and disease progression^10,11^. Rather, reduced mitochondrial function and energy production may also be important contributors to disease progression^10,12–15^. Mitochondrial structural changes and dysfunction have been widely described in MMA^12,16,17^. Previous liver and kidney tissue models have demonstrated MMUT-deficiency causes enlarged mitochondria and altered OXPHOS activity^16,17^. The consequences of this appear to be impaired mitophagy^12^ and shifted anaplerosis into the TCA cycle^10,18^. MMUT-KO cells and MMA patient fibroblasts have demonstrated a preference for reductive TCA cycle pathway, which was rescued by supplying cells with 2-dimethyloxoglutarate, an analogue of 2-oxoglutarate^10^. However, neither mitochondrial nor metabolic response to MMUT-deficiency in the CNS have been investigated due to the lack of an adequate model.

In the CNS, glutamine and glutamate have tissue-specific functions as stimulatory neuroactive molecules^19^. Additionally, mitochondria preferentially utilise OXPHOS for synaptic transmission and maintaining membrane potential through Na/K-ATPases^20–23^. Despite the clear role of neurological dysfunction in MMA and the long-term symptoms (i.e., metabolic stroke, epilepsy, optic neuropathy) invoked, no appropriate models have been employed to investigate this aspect of the disease. Such a shortfall is important because current treatments of MMA are symptomatic and do not protect against long-term CNS sequelae. Recent advances of patient-derived induced pluripotent stem cell (iPSC)-models offer an invaluable route to model neuronal dysfunction in MMA and provide valuable insights into CNS dysfunction.

In this study, we generated iPSC-derived neuronal models of MMA from variant-matched patient fibroblasts. We demonstrate that MMUT-deficient neurons have perturbed mitochondrial networks and reduced mitochondrial function. These appear to drive action potential attenuation in patient-derived neurons, potentially through reduced sodium currents. To tie metabolism to cellular function, metabolomic and transcriptomic exploration suggest altered glutamine processing, which is exacerbated by application of 2-dimethyloxoglutarate. Altogether, our results demonstrate that mitochondrial dysfunction may lead to a neuronal exhaustion-like phenotype, which may be an important component of neurological dysfunction in MMA.

## Results

### Generation of MMUT-deficient iPSC-derived neuronal lines

Deficiency of MMUT causes MMA, leading to biochemical alterations which drive disease pathology (Fig. 1a). Using Sendai virus based OKSM (OCT4, KLF4, SOX2, MYC) reprogramming, we generated iPSC clones from two unrelated individuals, each homozygous for the pathogenic, vitamin B_12_/cobalamin-unresponsive^24,25^ variant MMUT-p.(Asn219Tyr) (Pt1 & Pt2, *n* = 2; decimals indicate subclone, e.g. Pt1.1 is patient 1, subclone 1), and a wildtype control individual (Ct1, *n* = 1) (Fig. 1b and Supp. Fig. 1a). Isolated clones from all three iPSC lines did not retain Sendai virus markers used in ectopic pluripotency induction (Supp. Fig. 1b). These cell lines were characterised alongside a commercially available control (Ct2, *n* = 1). All four iPSC lines expressed the endogenous pluripotency markers: KLF-4, SOX2, OCT4 and NANOG, which were not found in the progenitor fibroblasts, except for KLF-4 which is natively expressed in fibroblasts (Supp. Fig. 1c). We demonstrated expression of the pluripotency markers NANOG and SSEA-4, and pan-stem cell marker SOX2, as well as the proliferative marker Ki67 (Fig. 1b), and could generate the three germ layers in all four cell lines (Fig. 1c). Comparative genomic hybridization indicated that induction of pluripotency did not generate *de novo* copy number variations between fibroblasts and their daughter iPSC clones (Supp. Fig. 2).

**Figure 1:**
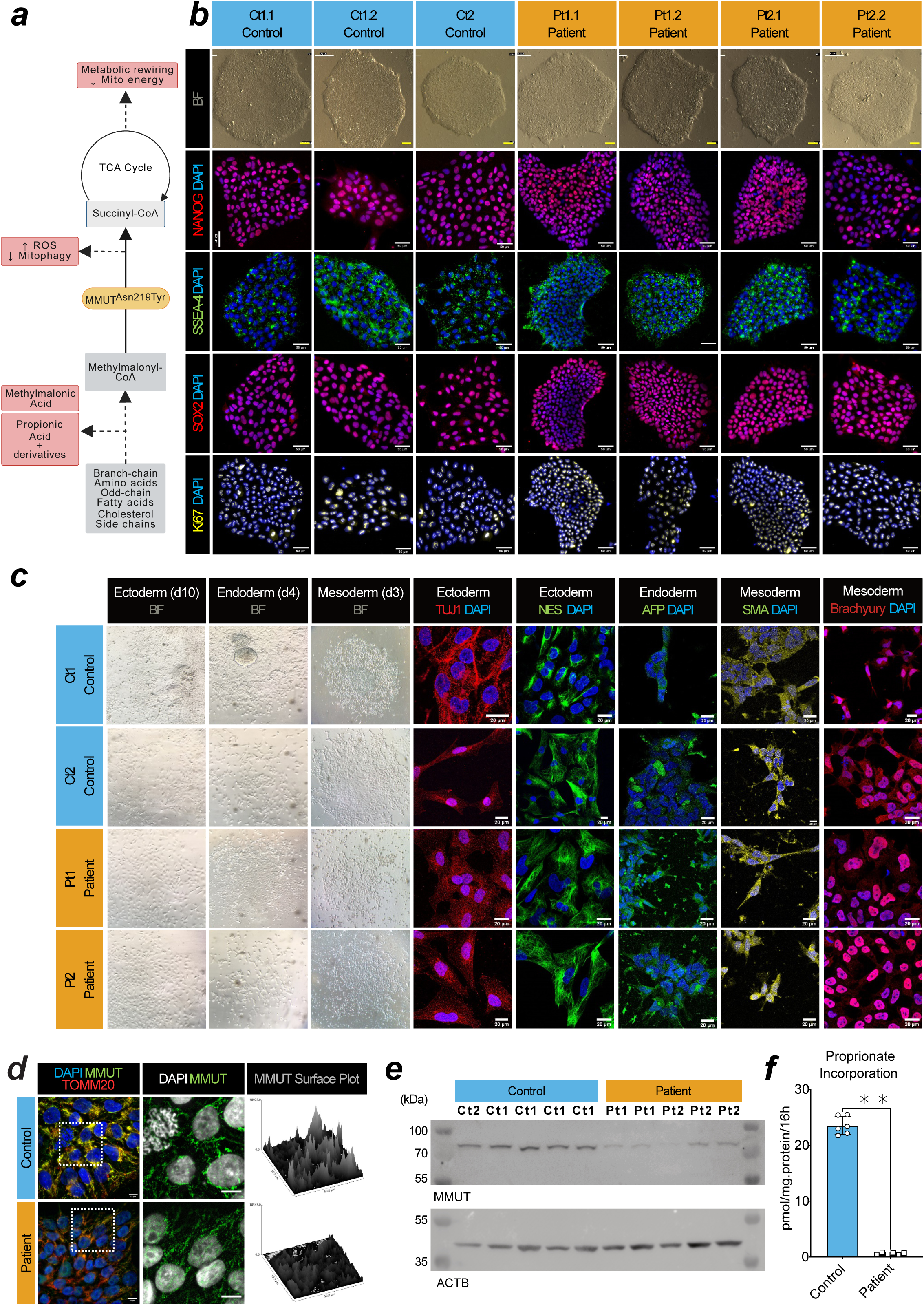
Generation of iPSC-derived MMUT-deficient neurons from individuals with methylmalonic aciduria. **a**, Schematic of the affected metabolic pathway in MMUT deficiency. Created in BioRender. **b,** Brightfield and epifluorescent acquisitions of pluripotency (NANOG, SSEA4, SOX2) and proliferation (Ki67) markers in representative iPSC cultures from control and patient lines. Scale, 50 μm. **c**, Ectoderm lineage indicated by beta-tubulin III (TUJ1) and nestin (NES) positive cells after 12 days *in vitro* (DIV). Endoderm lineage indicated by anti-α-Fetoprotein (AFP) positive cells after 4 days *in vitro* (DIV). Mesoderm lineage indicated by alpha-smooth muscle actin (SMA) and brachyury positive cells after 4 DIV. Scale, 10 μm. **d**, Comparison of the anti-MMUT (green) staining pattern from representative patient fibroblast-derived iPSCs compared to controls. Representative images come from cell line Ct1.2 and Pt1.1 For reference, anti-TOMM20, a mitochondrial protein, is also shown (red). Scale, 50 μm. **e**, Western blotting analysis of derived iPSCs using anti-MMUT. MMUT is anticipated at 83 kDa. The loading control ACTB is anticipated at 42 kDa. Uncropped membranes are available in Supp. Fig. 12. **f**, Propionate incorporation of two Ct1.1 and Ct1.2 wildtype iPSC sub-clones and Pt1 and Pt2 patient cell lines. Data points represent independent measurements taken from 3 separate cultures per cell line. Mean control iPSC incorporation values were 22.71 and 24.36 pmol/mg.protein/16h. Patient iPSC incorporation values were 0.77 pmol/mg.protein/16h in Pt1 and 0.44 pmol/mg.protein/16h in Pt2. P-value was tested using Mann-Whitney U-test (**, <0.01). Error is presented as standard deviation.

To evaluate whether the generated iPSC clones demonstrated MMUT deficiency, we examined MMUT protein expression by immunocytochemistry (Fig. 1d) and Western blotting analysis (Fig. 1e). Using a MMUT specific antibody (Supp. Fig. 1d and 1e), we observe a more punctate and less intense staining pattern, which does not co-localise well with translocase of the outer mitochondrial membrane complex subunit 20 (TOMM20) (Fig. 1d) and we found reduced MMUT protein in lysates of patient iPSCs (Fig. 1e), consistent with reduced enzymatic activity (2.58% of control), as demonstrated by an indirect activity assay which monitors propionate incorporation via MMUT into the TCA cycle^26^ (Fig. 1f).

To assess if MMUT deficiency affects neuronal development and function, we employed a 2D differentiation protocol that utilises dual-SMAD inhibition to generate dorsal forebrain-enriched neuronal progenitors^27,28^ (Fig. 2a). For all control and patient-derived iPSCs, we identified successful generation of neuroectodermal Nestin- and SOX2-positive neuroepithelium which progressively differentiated from a population including SOX2- and PAX6-positive neural stem cells (NSC) to TUBB3 and EOMES-positive neural progenitor cells and finally TUBB3, TBR1, MAP2, and NeuN-positive neurons (Fig. 2b). RT-qPCR revealed that patient cells express *SOX2*, *PAX6*, and *EOMES* at days 13 and 21 *in vitro* comparably to control cells when expression is normalized to day 0 (Fig. 2c). This suggests they adopt cortical NSC identity at the same rate as control cell lines. Immunocytochemistry (Fig. 2d) and Western blotting analysis (Fig. 2e) indicate MMUT protein expression is retained in NSC and neurons (Fig. 2d), however patient-derived neurons maintain the more punctate and less intense MMUT staining pattern and show reduced MMUT protein levels (Fig. 2e).

**Figure 2:**
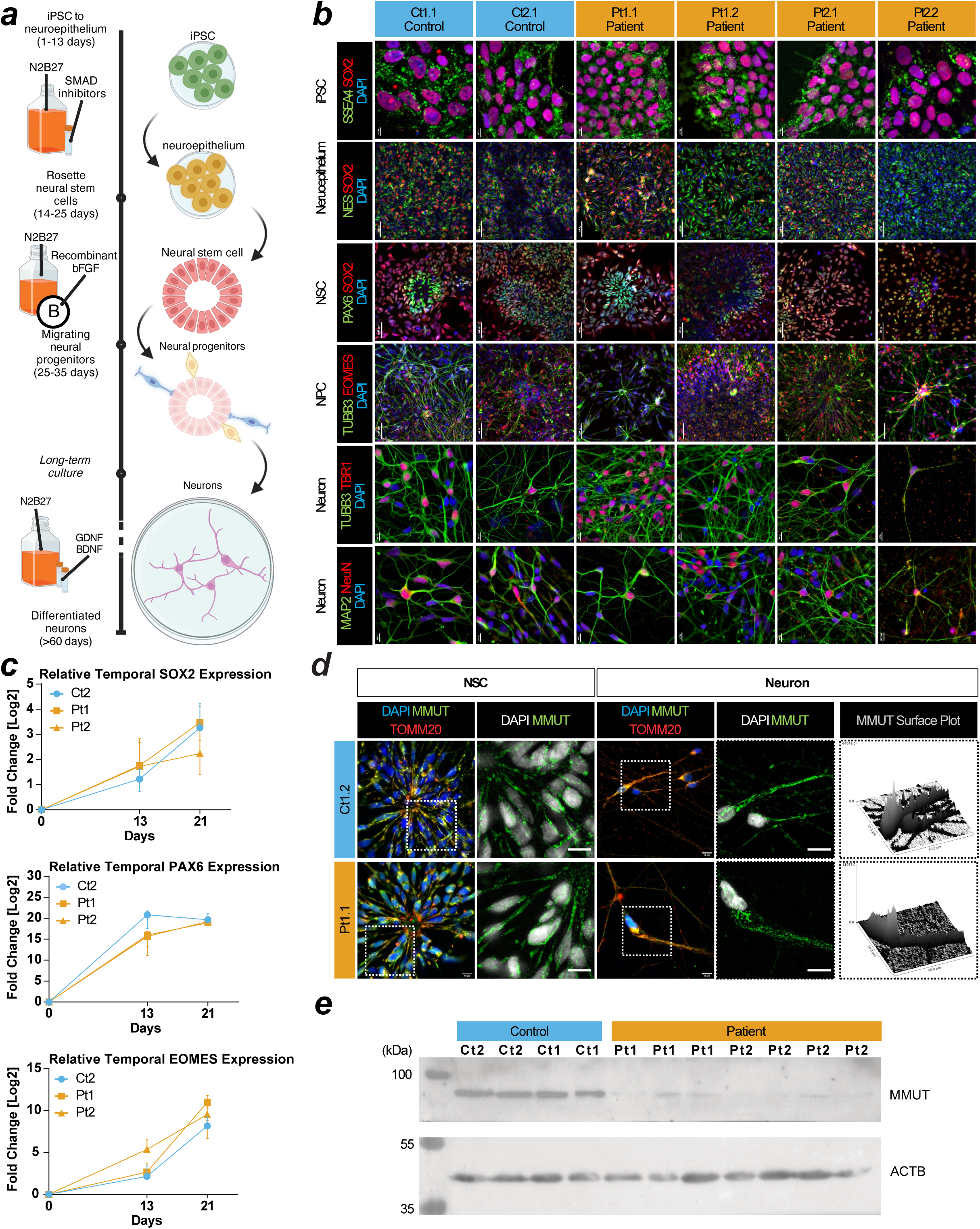
Derivation of cortical neurons from iPSCs affected by methylmalonic aciduria. **a**, Schematic of the 2D neuron differentiation protocol used in this article. Created using BioRender Forny, P. (2025) https://BioRender.com/w70y299. **b**, Staining for SSEA4 and SOX2 at day 0 for pluripotent stem cells (iPSC). Staining for Nestin (NES) and SOX2 at day 13 for neuroepithelium. Staining for PAX6 and SOX2 at day 21 for neural stem cells (NSCs). Staining for TUBB3 and EOMES at day 40 for cortical progenitors. Staining for TUBB3/TBR1 and MAP2/NeuN at day 50 signal postmitotic deep layer cortical neurons and pan-neuronal molecular markers in cultures, respectively. Scale: iPSCs and neurons have 10 µm bars. Neuroepithelium, NSC, and NPCs have 50 µm bars. **c**, RT-qPCR of SOX2, PAX6, and EOMES in iPSCs at day 0, neuroepithelium day 13, and NSCs day 21. Datapoints are representative of at least 3 independent experiments and error is reported as standard deviation. Expression values are relative to measurements at day 0 for each cell line. **d,** Immunocytochemistry of patient and control NSC and neurons. Scale, 10 μm. **e**, Western blotting analysis of neurons using anti-MMUT. MMUT is anticipated at 83 kDa. The loading control ACTB is anticipated at 42 kDa. Uncropped membranes are available in Supp. Fig. 12.

### Patient neurons have mitochondrial dysfunction and disrupted networks

TOMM20 is an essential mitochondrial protein often used as a proxy for mitochondrial abundance and shape^29^. Immunocytochemistry demonstrated altered TOMM20 staining patterns in patient-derived neurons that can be described as perinuclear and smaller (Supp. Fig. 3a), suggestive of disrupted mitochondrial networks. This was consistently observed in neurons derived from both patient lines, but not controls and was related to the extent to which MMUT could be detected within individual neurons (Fig. 3a and Supp. Fig. 3b). Investigation of spatial mitochondrial characteristics of patient neurons using TOMM20 staining revealed reduced mitochondrial volume and diameter but not sphericity (Supp. Fig. 3c,d). Further, disrupted mitochondrial networks in patient neurons appeared to impair mitochondrial response to OXPHOS insult, as demonstrated by comparable MitoSOX fluorescence at baseline but significantly reduced intensity after treatment with complex I inhibitor rotenone (Supp. Fig. 3e). In patient-derived cells, we identified two apparent populations, those expressing detectable MMUT (MMUT+) which overlapped with TOMM20, and those without (MMUT-) (Fig. 3a,b). To investigate the functional capacity of patient MMUT+ and MMUT− mitochondria, we used detectable fluorescence of tetramethylrhodamine-methyl-ester (TMRM) as a measure of mitochondrial membrane potential. We found a reduced TMRM signal in patient-derived neurons, a difference that was exacerbated in those cells with no detectable MMUT (Fig. 3c,d). We hypothesized that the lower TMRM signal denoted lower electron transport chain function, which we tested by inhibiting complex I with rotenone. We found reduced TMRM fluorescence in rotenone treated MMUT+ cells, less so in patient MMUT-, but not in control (Fig. 3d), suggesting that MMUT deficient mitochondria have a diminished proton motive force leaving them less able to react to events that would dissipate the membrane potential. We hypothesized that reduced membrane potential could trigger programmed cell death. Strikingly, this was supported by untreated patient-derived neurons having increased proportion of pro-apoptotic cleaved Caspase 3 (Fig. 3e) as well as DAPI nuclei staining patterns consistent with apoptotic cells (Fig. 3f and Supp. Fig. 3f).

**Figure 3:**
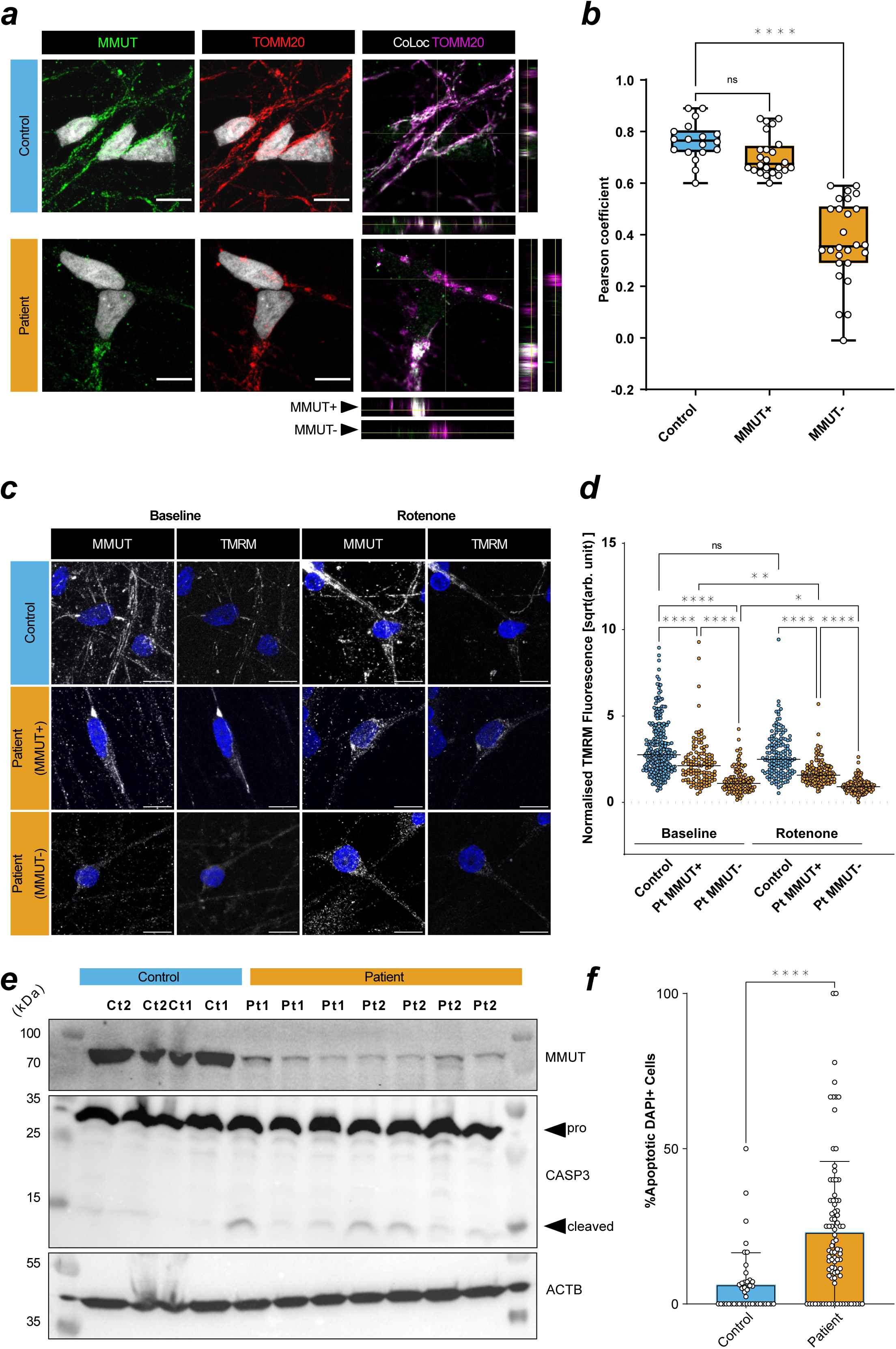
Patient neurons show mitochondrial dysfunction that overlaps with loss of MMUT protein. **a**, Left and middle: MMUT+ signal (green) and TOMM20+ mitochondrial signal (red). Right: TOMM20+ signal (magenta) and colocalization with MMUT (white). Representative images were selected from Ct1.2 and Pt1.2 cell lines. Orthogonal representative slices are also shown. Scale is 10 μm. **b**, Pearson correlation coefficient of TOMM20+ regions of interest (ROIs). One data point represents a Pearson’s coefficient from one ROI. 3-5 ROIs per image, total images used from control *n* = 6 (18 total ROIs) and from patient *n* = 11 (50 total ROIs). P-value determined by one-way ANOVA with multiple testing (ns, >0.05; ****, <0.0001) Datapoints represent Ct1.2, Ct2, Pt1.1, Pt1.2, and Pt2.1. Error is plotted as min. to max. **c**, Neuronal mitochondria stained with MMUT and TMRM (both grey). Scale is 10 μm. Representative images were selected from Ct2, Pt1.1, and Pt2.1 **d**, TMRM+ ROIs selected from non-overlapping confocal images. Cumulative data is representative of *n* = 6 in control without rotenone, *n* = 4 in control with rotenone, *n* = 8 in patient without rotenone, *n* = 9 in patient with rotenone. One datapoint represents one background/size-corrected ROI. P-value determined by Kruskal-Wallis with multiple testing (ns, >0.05; *, <0.05; ****, <0.0001). Datapoints represent Ct2, Pt1.1, and Pt2.1. Bar represents the median value. **e**, Western blotting analysis of MMUT, caspase-3 (CASP3), its cleaved product, and ß-actin (ACTB) in total cell lysate from untreated neurons. Each column represents an independent lysate. Uncropped membranes are available in Supp. Fig. 12. **f**, Quantification of DAPI+ apoptotic cells determined from representative, non-overlapping, epifluorescent images acquired using a 63x objective. Images were taken of fixed untreated neurons at day 50 *in vitro.* Datapoints in control represent Ct1.2 and Ct2, and datapoint in patient represent Pt1.1, Pt1.2, Pt2.1, and Pt2.2. Cumulative data is representative of *n* = 43 (776 cells) in control and *n* = 87 (1317 cells) in patient images. Significance is assessed by Mann-Whitney test (****, <0.0001). Error is standard deviation. Dunn’s multiple comparisons test used in b, d.

### Electrophysiological assessment reveals an exhaustion phenotype in patient-derived neurons

In order to assess generated neurons’ functional properties, we performed whole-cell analysis on patient (*n* = 132, Pt1 & Pt2) and control (*n* = 94, Ct1 & Ct2) neurons (Supp. Tab. 1). Similar to previous reports^30^, principle component analysis (PCA) of all 31 extracted features from step current injection protocols (Supp. Tab. 2) revealed three distinct clusters (Supp. Fig. 4a), which were further discriminated by utilizing 21 primary variables (Fig. 4a and Supp. Fig. 4b). These clusters could be best represented as non-action potential (AP) firing (Type 1, T1), single-AP firing (T2), and multiple AP-firing (T3) neurons (Fig. 4a,b and Supp. Fig. 4a). They were not separated by genotype (patient vs control) (Supp. Fig. 4b), but rather by variation within inherent electrophysiological properties, such as resting membrane potential, input resistance, capacitance, number of APs, and AP threshold (Supp. Fig. 4c). Unguided hierarchical clustering of 21 primary variables separated assessed neurons by T1, T2 or T3, but not by genotype (Supp. Fig. 4d). However, we observed an apparent decrease in T3 and corresponding increase in T2 in patient-derived neurons (Fig. 4c), suggesting that generating successive APs may be corrupted in patient neurons.

**Figure 4:**
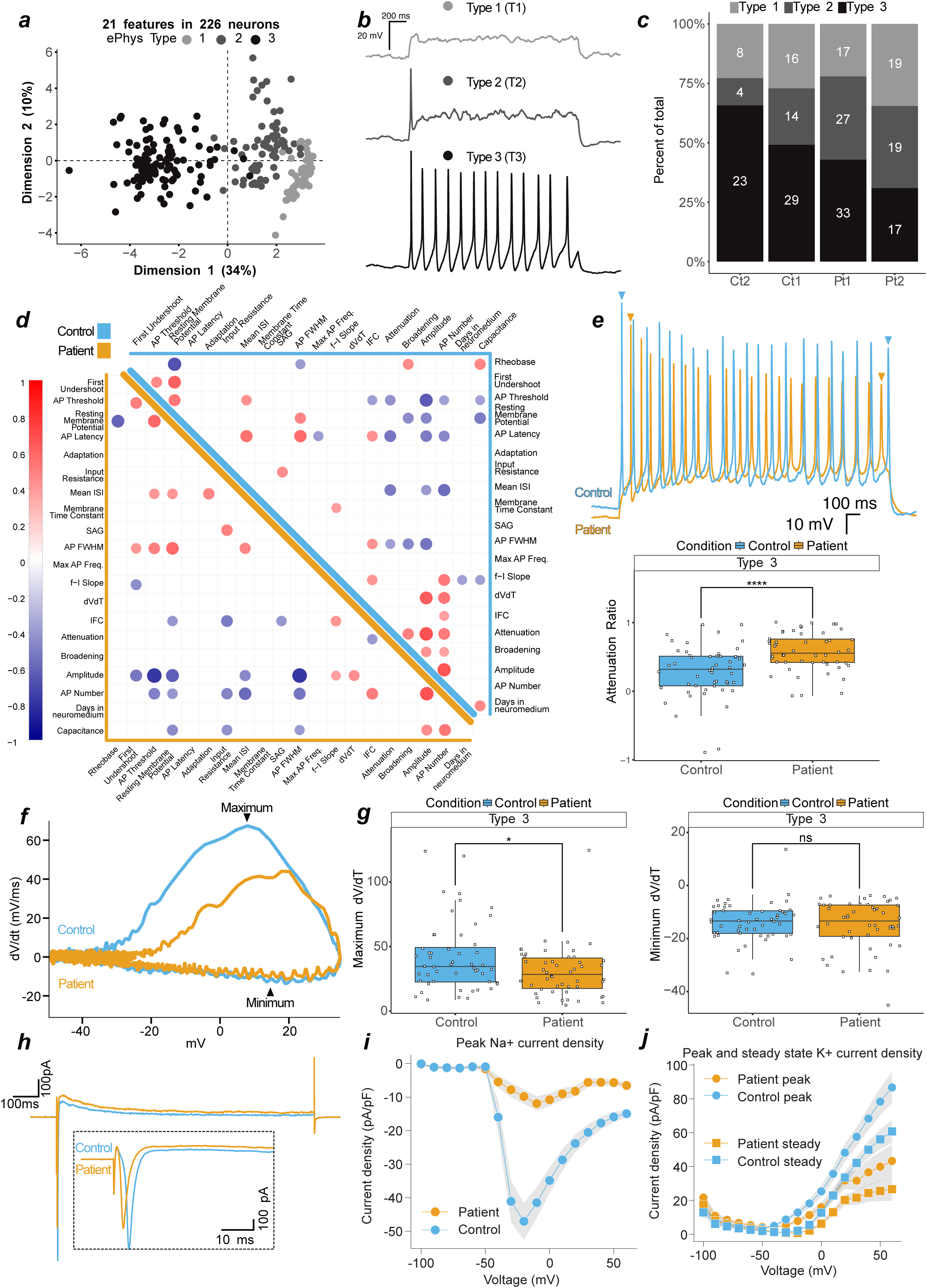
Patient neurons show action potential exhaustion driven by reduced sodium currents. **a**, First two dimensions (44% variance explained) of a principal component analysis of 21 measured features from patient (*n* = 132 (Pt1 n=77, Pt2 n=55)) and control (*n* = 94 (Ct1 n=59, Ct2 n=35)) neurons. Greyscale represents 3 identified neuronal types, type 1 (light grey, T1, *n* = 60), type 2 (dark grey, T2, *n* = 64), and type 3 (black, T3, *n* = 102). **b**, Current-clamp, representative traces from T1, T2, or T3 neurons in response to depolarizing current injection. **c**, Proportional generation of classified neurons of control (Ct1, Ct2) and patient (Pt1, Pt2). **d**, Correlation matrix of 21 features in patient (bottom-left, orange) and control (top-right, blue) neurons. Positive correlations indicated by red and negative by blue. Features with significance *P* = <0.01 are displayed on the matrix, values above are blank. **e**, Top, overlayed traces from T3 control and patient neurons. Bottom, attenuation ratio from 102 measured T3 neurons. **f**, Phase plots of 2 overlayed single action potentials from representative T3 control and patient neurons. **g**, Depolarisation (left, maximum dV/dt) and repolarisation (right, minimum dV/dt) velocities. **h**, Voltage-clamp, representative overlay of currents evoked by step to −30 or −20 mV in control and patient neurons, respectively. Inset is an enlarged section of the initial segment. **i**, Peak control (*n* = 67) and patient (*n* = 66) sodium current densities. **j**, Peak and steady control (*n* = 67) and patient (*n* = 66) potassium current densities. In i-j, current densities are normalised to capacitance, and variation is SD and present as shaded area. In e and g, datapoints represent one neuron, data are pooled into control (2 independent lines) and patient (2 independent lines), significance is by t-test and reported with p-value adjustment using Holm, and whiskers represent 1.5·IQR.

To explore this further, we performed correlation analysis on 21 primary variables in T3 neurons derived from patients (*n* = 50) and controls (*n* = 52) (Fig. 4d). For all cells, we identified anticipated correlations between variables, such as AP threshold and AP full width at half maximum (FWHM, i.e. time between slope rise and fall at half-max amplitude) with AP amplitude (negative correlations), as well as AP amplitude with AP number and resting membrane potential with AP FWHM (positive correlations) (Fig. 4d and Supp. Fig. 5a,b). Striking, however, was that variables expected to correlate with AP attenuation, and found to do so in control neurons (e.g., AP amplitude, AP threshold, and FWHM), did not correlate in patient neurons (Fig. 4d and Supp. Fig. 5c). Closer inspection of the reduction of AP amplitude following multiple APs (measured as the attenuation ratio), revealed a significantly stronger loss of AP amplitude and hence enhanced attenuation in patient neurons (Fig. 4e). This is indicative of a mitochondrial exhaustion-like phenotype, which is mirrored in Leigh Syndrome models that lacked repetitive spiking and contained higher numbers of non-spiking neurons^31^.

To link the loss of AP amplitude over multiple firings and mitochondrial dysfunction, we further assessed attenuation in selected cells whose mitochondrial morphology was visualized using MitoTracker™ (Supp. Fig. 5d). Consistent with previous data (Supp. Fig. 3a-d), all control cells appeared to have intact mitochondrial networks, whereas patient-derived cells had either intact or disrupted mitochondrial networks (Supp. Fig. 5d). Patient cells with intact mitochondrial networks had an attenuation ratio similar to control neurons, whereas neurons with disrupted mitochondrial networks had increased attenuation (Supp. Fig. 5e), suggesting an association between AP exhaustion and mitochondrial dysfunction.

Depolarisations generating higher AP amplitudes are achieved by influxes of cations; the velocity of these influxes can be calculated through the first derivative of the membrane potential slope over time (dVdt, Fig. 4f). dVdt ratios in patient-derived neurons were decreased compared to controls (Supp. Fig. 5f), which was driven by the depolarisation (reported as maximum) and not repolarisation (potassium efflux, reported as minimum) phase (Fig. 4g). In control neurons, the maximum dVdt positively correlated with the attenuation ratio, a relationship which was lost in patient-derived neurons (Supp. Fig. 5g). Similarly, the positive correlation of dVdt with AP number and with AP amplitude found in control neurons was lost or reduced in patient-derived cells (Fig. 4d and Supp. Fig. 5h). These findings suggest that patient-derived cells cannot sustain sufficient current densities to maintain action potential firing, which may lead to the observed exhaustion-like phenotype.

To test the hypothesis that reduced sodium currents drive these observations, we performed additional voltage-clamp recordings on patient (*n* = 92) and control (*n* = 89) neurons. The selected neurons contained the same patient T3 proportional decrease (Supp. Fig. 6a), increased attenuation and decreased dVdt ratio as the overall population (Supp. Fig. 6b). We found substantially reduced sodium current densities in patient-derived neurons when compared to control, with peak densities reduced 5-fold in patient-derived neurons (Fig. 4i,j; Supp. Fig. 6c-e).

### Rewired glutamine-glutamate metabolism alters glutamatergic identity

We next utilised targeted metabolomics to investigate the connection between mitochondrial dysfunction and altered neuronal excitability. MMA related metabolites, including hydroxypropionic acid and propionyl-carnitine were elevated as expected in patient-derived neurons, while lactate tended to be increased with methylmalonic acid and 2-methylcitrate appearing unchanged (Fig. 5a and Supp. Fig. 7a). Despite our finding of disturbed mitochondrial networks in patient-derived neurons, we found no difference in the pools of the anaplerotic intermediates pyruvate, alanine, glutamine and glutamate (Supp. Fig. 7b) as well as TCA cycle intermediates (Supp. Fig. 7c) between patient and control-derived neurons, which is consistent with findings in patient fibroblasts^10^. This prompted us to investigate the anaplerotic contribution to the TCA cycle from labelled glutamine (Fig. 5b). Glutamine flows into the TCA cycle via glutamate and 2-oxoglutarate (Fig. 5b). Following labelled glutamine supplementation, we found a high proportion of labelled glutamine, which was reduced in glutamate and 2-oxoglutarate, but found no difference between fractional labelling (Supp. Fig. 7d) nor pool levels (Supp. Fig. 7e) of all three intermediates between patient and control-derived neurons. We also found no difference in the fractional incorporation nor pool levels of measured TCA intermediates (Supp. Fig. 7f,g). However, we did notice reduced fractional incorporation, but not pool levels, of aspartate (Fig. 5c). Glutamine-derived aspartate can be produced via oxaloacetate by cytosolic or mitochondrial aminotransferases, whereby the direction of the TCA cycle (oxidative or reductive) influences the number of labelled carbons (Fig. 5b). We found the ratio of M+5 to M+4 citrate, and M+3 to M+4 malate (M+*x* refers to the number of carbon isotopes that are present, where a ratio of 1 indicates equal contribution from oxidative and reductive), to be changed in patient-derived neurons (Fig. 5d). This altered ratio is supportive of metabolic rewiring in the sense that labelled carbons entering the TCA cycle are affected in patient-derived cells, which is also supported by altered aspartate labelling.

**Figure 5:**
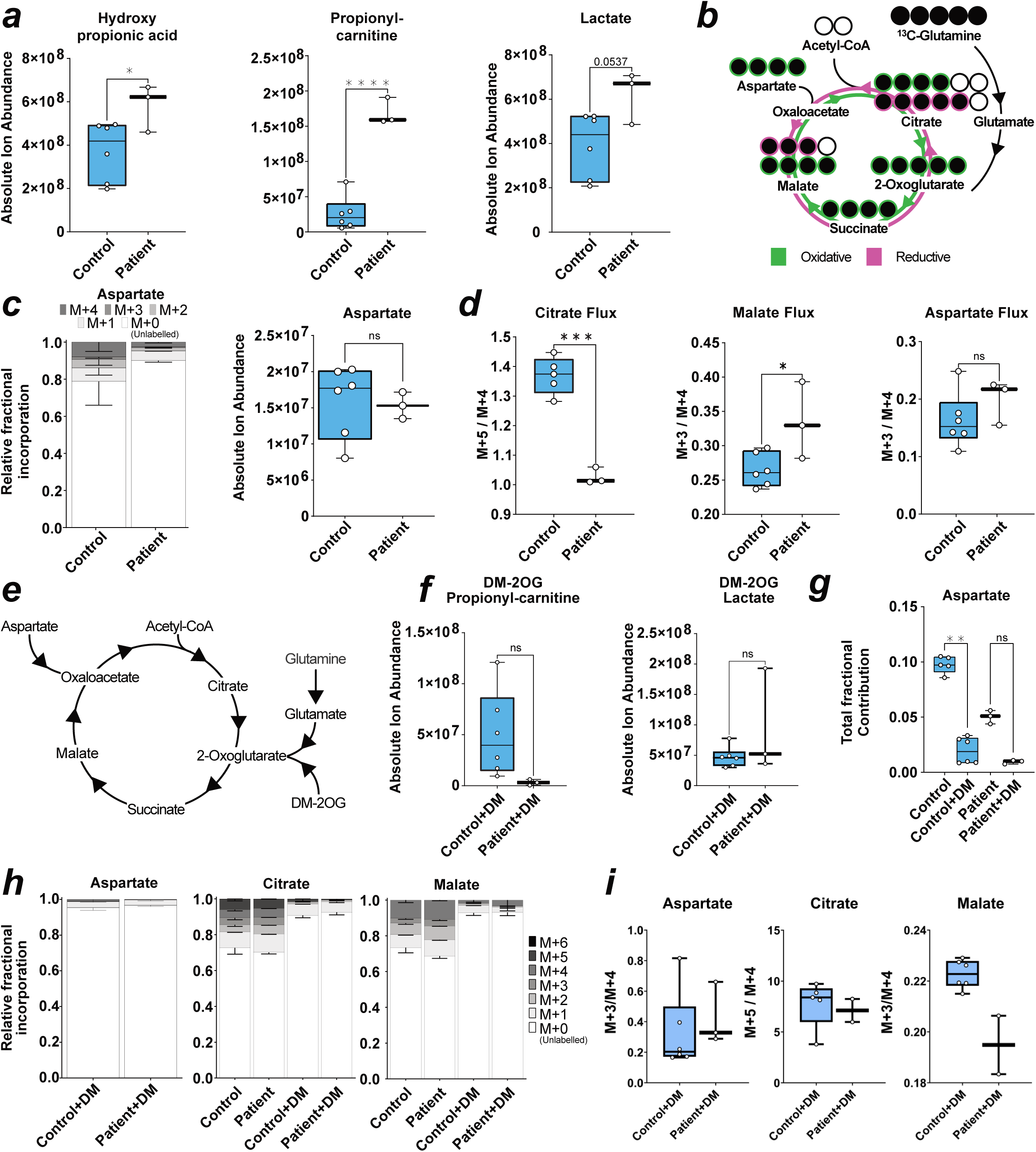
Metabolomics reveal dysregulated glutamate and glutamine neuronal metabolism. **a,** Box plots of absolute ion abundance of disease-related metabolites. P-value is reported as one-way ANOVA for hydroxypropionic acid, and unpaired t-test for propionyl-carnitine, and lactate. **b**, Schematic of labelled ^13^C-glutamine metabolism into the TCA cycle. Labelled carbons = filled black circles. **c** Stacked bar chart of contribution of labelled glutamine carbons to aspartate (left) and box plot of absolute ion abundance of aspartate pool (right). P-value calculated via unpaired t-test. **d**, Box plot of ratio between M+5 to M+4 labelled citrate fractions (left) and box plot of ratio between M+3 to M+4 labelled malate or aspartate fractions (right). P-value is reported as unpaired t test. **e**, Representative schematic for the entry of dimethyl-2-oxoglutarate (DM-2OG) into the TCA cycle and selected anaplerotic entry points. **f**, Box plot of absolute ion abundance of propionyl-carnitine and lactate in DM-2OG treated neurons. *P* value is reported as unpaired t test. **g**, Box plot of total fractional contribution of carbons to untreated and DM-2OG treated neuronal abundance of aspartate. P-value is reported as Mann-Whitney (left). **h,** Stacked bar chart of contribution of labelled carbons to untreated and DM-2OG treated neuronal abundance of malate (left) and citrate (right). **i,** Box plots of ratio between M+3 to M+4 labelled aspartate and malate fractions (left and right) in DM-2OG treated neurons and box plot of ratio between M+5 to M+4 labelled citrate fractions (middle) in DM-2OG treated neurons. In **a, c, d, f, g, h,** error = min/max (box plot) or SD (fractional bar plot). Each datapoint represents 3 technical replicates from one independent sample. Genotype conditions are pooled into control (blue) and patient (no colour). In significance tests: ns = >0.05, * = <0.05, *** = <0.001, **** = <0.0001. In all panels, control is represented by Ct1.2 and Ct2. Patient is represented by Pt1.2 and Pt2.2.

We also investigated the pool sizes of glycolytic intermediates, which follows report of increased hexoses in primary patient fibroblasts^10^. Using a targeted approach to measure levels of specific detectable sugar phosphates^32^, we identified elevated galactose-1-phosphate, mannose-6-phosphate, and glucosamine-6-phosphate in patient-derived neurons compared to controls (Supp. Fig. 8a). We found reduced *N*-acetyl-glucosamine-6-phosphate and *N*-acetyl-mannosamine-6-phosphate (Supp. Fig. 8b) and elevated nucleotide sugars GDP-fucose and GDP-mannose (Supp. Fig. 8c). We suspect this may be indicative of a reactive shift to glycolysis, as OXPHOS in disease contexts is impaired, which is a phenomena also described in hypoxia^33^. We found no change in other glucose or fructose phosphate derivatives nor pentose phosphate derivates (Supp. Fig. 8a,d and Supp. Tab. 3). Our finding of elevated galactose-1-phosphate and mannose-6-phosphate suggest glycolytic flux is increased. Further investigation is required to determine a link between increased glycolytic metabolites and rewired TCA cycle metabolism, as this phenomena has been reported previously^10^.

Based on the above changes, we attempted to rescue the phenotype in patient-derived neurons using dimethyl-2-oxoglutarate (DM-2OG) a membrane permeable analogue of the TCA cycle intermediate 2-oxoglutarate (Fig. 5e). Application of DM-2OG increased some but not all pools of TCA cycle intermediates measured, including glutamate and glutamine (Supp. Fig. 9a-b), and resulted in amelioration of the difference between pool levels of hydroxypropionic acid, propionyl-carnitine, and lactate (Fig. 5f, Supp. Fig. 9c) as well as fractional contribution into aspartate of control and patient-derived neurons and normalisation of TCA cycle flux (Fig. 5g-i). DM-2OG supplementation increased selected hexose phosphates but patient galactose-1-phosphate, mannose-6-phosphate, and glucosamine-6-phosphate remained significantly increased compared to treated control neurons (Supp. Fig. 9d). However, DM-2OG treatment reduced the difference between patient and control *N*-acetyl-glucosamine-6-phosphate and *N*-acetyl-mannosamine-6-phosphate (Supp. Fig. 9e), which we attribute to DM-2OG flooding the TCA cycle and conversion of glutamine to glucosamine that then inhibits side pathways^34^.

### TCA cycle rewiring impacts neuronal excitability at the synaptic level

DM-2OG altered the neuronal levels of neurotransmitter amines, such as glutamate, leading us to investigate the effect of DM-2OG supplementation and subsequent metabolic rewiring on electrophysiological properties. We performed current-clamp recordings on 36 neurons with supplementation of DM-2OG between 0.1 – 6.0 mM (Supp. Fig. 10a). We found that higher doses abolished AP firing capacity (Supp. Fig. 10a), which may be linked to increasing hyperpolarisation and decreasing capacitance (Supp. Fig. 10b-c). Voltage-clamp recordings on control neurons demonstrated that 1 mM and 6 mM DM-2OG reduced sodium current densities to levels that were similar to untreated patient-derived neurons (Fig. 6a,b). At lower doses, control but not patient-derived neurons were still able to produce APs and maintain sodium current densities (Fig. 6a-b and Supp. Fig. 10a). We hypothesized that patient metabolic rewiring altered neuronal function at the synaptic level. We investigated the packaging of glutamate in the synapse through *SLC17A7*, which encodes the vesicular glutamate transporter 1 (VGLUT1). We found a greater number of VGLUT1+ foci in patient neurons (Supp. Fig. 10d,e), suggesting a predisposition towards glutamate excitotoxicity that could contribute to the exhaustion phenotype, and, more generally, to the neurological phenotype. Consequently, we recorded spontaneous excitatory postsynaptic currents of untreated patient (*n* = 46) and control (*n* = 48) as well as 0.1mM DM-2OG treated patient (*n* = 7) and control (*n* = 8) neuronal networks. We observed an increased charge of synaptic events in patient-derived neurons (Fig. 6c,d). However, we did not observe difference in event amplitude, decay kinetics, or frequency (Supp. Fig. 10f-h). The event duration (charge) may be related to glutamate receptor subunits, which combine to form functional neuronal glutamate receptors^35^.

**Figure 6:**
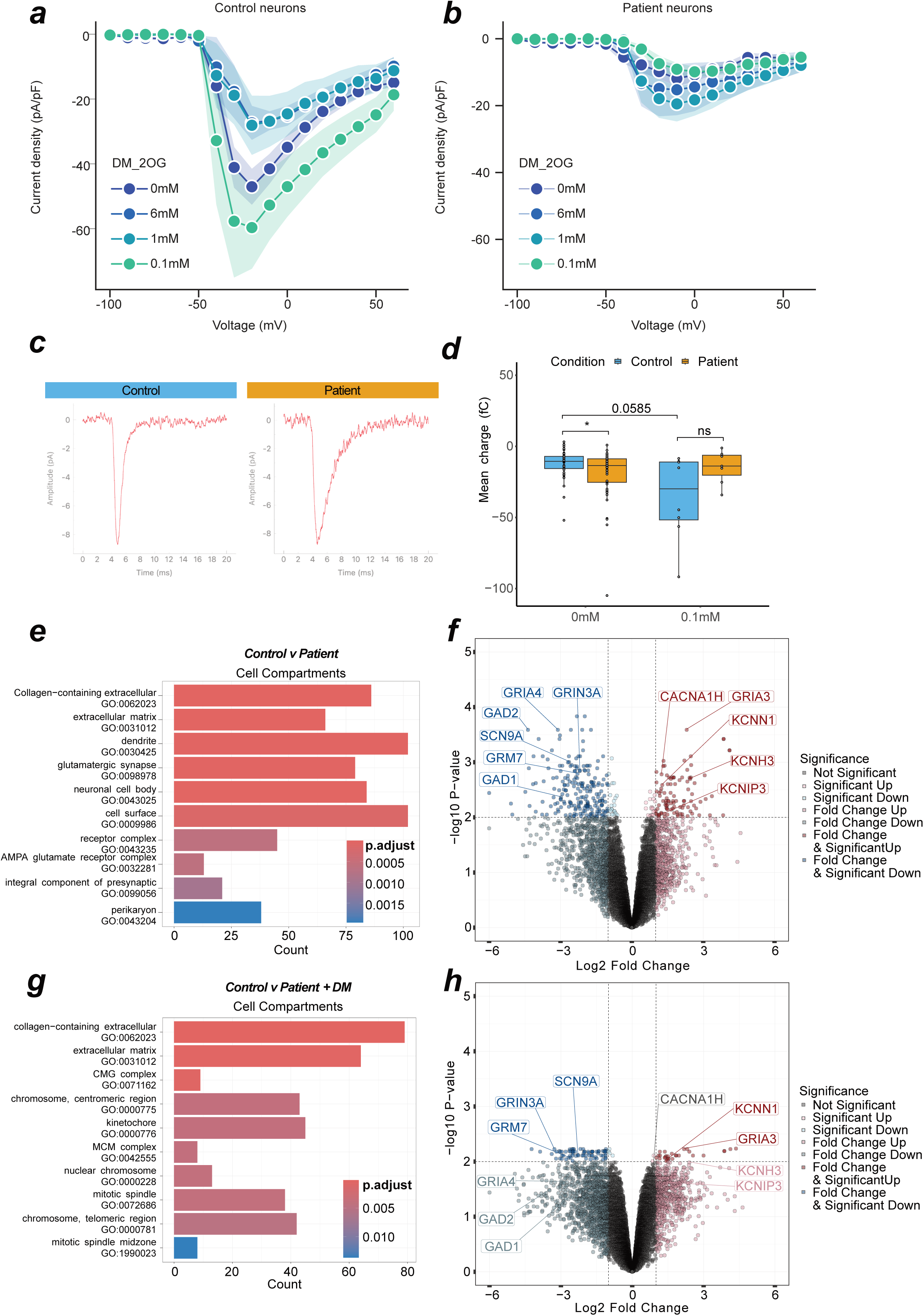
Metabolic rewiring affects glutamatergic synaptic transmission. **a,** Line graph indicating maximum current density at evoked voltages in control neurons with various DM-2OG treatments. Datapoint represents at least 5 independent samples, error = SD represented by shading. **b,** Line graph indicating maximum current density at evoked voltages in patient neurons with various DM-2OG treatments. Datapoint represents at least 6 independent samples, error = SD represented by shading. **c,** Representative averaged traces of spontaneous synaptic currents from single control (n events = 114) or patient (n events = 36) neurons in voltage-clamp. **d,** Box plot of spontaneous synaptic event charge of individual neurons in voltage-clamp. *P* value is calculated via mann-whitney **e,** Over-representation analysis of top 10 up and down-regulated GO terms from untreated control (*n* = 5) and patient (*n* = 6) derived neurons. Count refers to number of contributing genes. **f,** Volcano plot of differential regulation between untreated control (*n* = 5) and patient (*n* = 6) neurons. **g,** Over-representation analysis of top 10 up- and down-regulated GO terms. Count refers to number of contributing genes from untreated control (*n* = 5) and 0.1 mM DM-2OG treated patient (*n* = 4) derived neurons. **h,** Volcano plot of differential regulation between untreated control (*n* = 5) and 0.1mM DM-2OG treated patient (*n* = 4) neurons. In **c, d,** each datapoint represents one independent sample. Genotypes are control (blue) and patient (orange). In **a, b** center is the mean and error are SD. In **c, e,** colour represents *P* value magnitude. In **f, h,** values are FDR, genes highlighted contribute to a top differentially regulated GO term. Dotted lines indicate Log2 fold change of 1 and −log10 P-value of 0.01. In **d,** box plot is median and min/max. In panels a-d, control is represented by Ct1 and Ct2, and patient is represented by Pt1 and Pt2. In panels e-h, control is represented by Ct1.2 and Ct2, and patient is represented by Pt1.1, Pt1.2, Pt2.1, and Pt2.2.

Given the electrophysiological and metabolic changes in patient-derived neurons, we performed bulk transcriptomics on control and patient neurons. Analysis of the neuronal transcriptome via PCA revealed successful separation of controls and patients based on inherent variance (Supp. Fig. 11a). Using over-representation analysis, we found gene networks involved in “glutamatergic synapse”, “ion channel activity”, and “chemical synaptic transmission” to be differentially expressed (Fig. 6e and Supp. Fig. 11b-c). Further analysis revealed glutamate packaging machinery (*SLC17A7*), and glutamate receptors, such as *GRM7*, *GRIN3A*, *GRIA3*, *GRIA4*, to have altered expression in patient-derived neurons (Fig. 6f and Supp. Fig. 11d). We also observed several ion channel genes, including sodium (*SCN9A*), calcium (*CACNA1H*), and potassium (*KCNH3*, *KCNN1*, *KCNIP3*) channels, to be differentially expressed (Fig. 6f). Moreover, GAD1/2, which catalyses the production of 4-aminobutanoate (GABA) from glutamate, depending on the supply pool size of glutamate in the cytosol, appeared to be downregulated in patient-derived neurons (Fig. 6f).

To test if subunit composition is altered, we treated control and patient neurons with 0.1 mM DM-2OG and analysed the transcriptome. Comparison of untreated control and DM-2OG supplemented patient-derived neurons revealed the absence of many previously identified significantly differentially regulated synaptic related GO terms (Fig. 6g). Of previously identified glutamatergic synapse differentially expressed genes, only GRM7, GRIN3A, SCN9A, KCNN1, and GRIA3 were still significant (Fig. 6h). Disease-state metabolic rewiring changes the TCA cycle, an impact that we can replicate with DM-2OG treatment. We demonstrate DM-2OG increases metabolite pools, such as glutamate, and these treatments recapitulate aspects of the exhaustion phenotype.

Finally, we leveraged our bulk transcriptomic data to perform RNA-seq deconvolution protocols to achieve pseudo single-cell RNA-seq data for cell type. This analysis revealed that the largest cellular population across all cell lines were those with excitatory neuron cellular identity (Supp. Fig. 11e), although each population was heterogenous and also contained inhibitory neurons and astroglia precursors, in line with the findings of the original methods paper^27^. We were also able to extract specific gene transcripts that identify specific cellular populations such as region-specific proliferating cells (PCNA, DCX, PAX3, PAX6, HES6), discrete neuronal populations (SLC17A6, GAD2), and astrocytic precursor populations (GJA1, ADGRV1) (Supp. Fig. 11f). DM-2OG was not found to have any consistent effect on cell type across the various lines (Supp. Fig. 11e). As patient liver biopsy had found reduced protein expression of cytochrome c oxidase (COX)^36^, we report mitochondrial COX transcripts (MT-CO1, MT-CO2, MT-CO3) are not differentially expressed, conversely subunits COX5A and COX6B were upregulated in patient-derived neurons (Supp. Fig. 11g).

## Discussion

In this study, we have gained insight into the conserved role of mitochondrial disruption in MMA and uncovered an exhaustion-like phenotype in neurons affected by mitochondrial disease. We described how mitochondrial disruption and reduced sodium current density underlie AP attenuation. MMUT deficiency rewires neuronal metabolism and conferred a greater demand of glutamine and aspartate to the TCA cycle, we also documented reduced current density through modulation of synaptic ion channel dynamics. We introduce the first *in vitro* human-relevant neurological model of MMA. We were able to model one common European pathogenic variant c.655A>T. However, we were unable to generate an isogenic control due to the genomic landscape of the mutation site. A further caveat is the use of only two control lines, and a lack of neural cell-type characterization using single cell RNA-seq. However, since we observed comparable rates of cortical marker adoption and gross electrophysiological functionality of patient-derived cells were indistinguishable from control neurons, we expect that the excitatory populations generated across conditions were comparable. Nevertheless, we make the final note that experiments making use of MitoTracker and TMRM were not performed with an additional cell-type specific maker, and we therefore cannot determine exactly which cell-types are most prone to mitochondrial dysfunction.

The occurrence of neurological symptoms despite treatment highlights the need to explore neuronal pathomechanisms^37–43^. Our findings reinforce previous comprehensive evidence that mitochondrial dysfunction drives pathophysiology in MMA^12,16,17^. These evidence fit well within the current literature that describe reduced mitochondrial respiration and/or mitochondrial ultrastructure alterations in primary patient kidney cells^12^; liver, kidney, heart, and skeletal muscle of patients^13^; Mut^−/−^ mice kidney^17^ and liver^16^. Here we demonstrate that mitochondrial dysfunction is also an important component of neuronal pathology. Although electrophysiology has never been assessed in MMA, two separate iPSC-derived models of the primary mitochondrial disease Leigh Syndrome demonstrate that pathogenic variants in the electron transport chain reduced oxygen consumption^31,44^. Electrophysiological reporting was mixed, one study found SURF1 mutations reduced sodium and potassium currents, which resulted in reduced repetitive spiking^31^. Another paper concluded that no difference between patient and control cells’ intrinsic properties were found^44^.

Although the term “attenuation” is not used in these referenced papers, it is clear that reduced sodium currents and decreased AP amplitudes are present^31^ and current-evoked AP amplitudes decrease over time. This phenomenon is demonstrated more clearly in a current ramp test, however both examples demonstrate an exhaustive phenotype comparable to attenuation^44^. These findings can be related back to biophysical studies in sodium inactivation which posit that complex gating can be tied to the specific subpopulation of sodium channel by the incorporation of specific subunits^45^. This consideration is even more striking when we consider our findings of differential regulation of a number of sodium channel subunits.

Neurons have been shown to have a greater reliance on OXPHOS during AP firing^46,47^. One route to meeting such a demand in the brain is through the oxidation of glutamine^48–51^, which is coupled to incremental glucose oxidation^51^. Derangement of OXPHOS puts a greater reliance on glutamine and aspartate metabolism^52^. Our finding of reduced glutamine contribution to aspartate suggests rewiring of the TCA flux^53^, consistent with reduced reliance on reductive TCA cycle flux in patient-derived neurons. This pattern is consistent with a greater importance on anaplerotic metabolites. Elevated levels of lactate and hexose phosphate sugars suggest glucose flowing into glycolysis may also be rewired to side pathways^54^. Altered dependence of patient neurons on glutamine and aspartate may feasibly lead to differential expression of glutamatergic receptor subunits. Hippocampal neurons rapidly redistribute their glutamate receptors after glutamate and glycine treatment^55,56^, indicating that metabolic shift may be strong enough to cause the differential expression of ionotropic glutamate receptors observed in patient neurons. Concurrently this may result in patient neurons’ prolonged synaptic events or reduced ability to sustain successive action potentials. *GRIA4*, which is downregulated in patient cells, is the fastest glutamate receptor and its expression is linked to receptor kinetics^57^. GRIA4-containing AMPA receptors also appear responsive to changes in spontaneous synaptic events^58^.

Previous work in our laboratory found disease-specific biochemical alterations in patient fibroblasts^10^. Crucially, our findings mirror the results from patient fibroblasts in a reduced propionate incorporation, elevated propionyl-carnitine and hydroxy-propionate, although we did not identify increased methylmalonic acid nor 2-methylcitrate. Manipulation of disease-state metabolic rewiring with DM-2OG demonstrated a rescue of disease-related metabolites but exacerbated components of the exhaustion phenotype in control-derived neurons. This suggests an association between metabolic rewiring and gene expression. Decreased measurements of *N*-acetyl-glucosamine-6-phosphate and *N*-acetyl-mannosamine-6-phosphate may link changes in the hexosamine biosynthesis pathway to altered transcription, as this pathway contributes posttranslational modifications to intracellular proteins, including glycosylation and associated transcription factors. Furthermore, the use of DM-2OG and its effect on glutamate, a neurotransmitter, evidenced these effects by abolition of AP firing capacity.

To date, great focus has been placed on the basal ganglia, due to the occurrence of bilateral lesions within the basal ganglia, we chose to focus our efforts on cortical neurons. The reasoning behind this decision was three-fold: (1) protocols to generate glutamatergic neurons of the cortical forebrain are better characterised and more sophisticated than GABAergic neurons or cholinergic interneurons of the telencephalon. (2) In individuals affected by MMA, bilateral lacunar infarctions of the basal ganglia are often reported after acute metabolic decompensation, which produces movement disorder, however other neurological symptoms such as delayed myelination, subcortical white matter changes, cortical atrophy, and brain stem and cerebellar changes are also commonly reported. From this list of symptoms, multiple cell types are affected, however no information has been published concerning which cellular populations generate these symptoms mechanistically. Moreover, whilst we and others have shown that MMUT is expressed in neurons^59^, it has yet to be demonstrated to be expressed in astroglia, microglia, or oligodendrocytes, and yet is believed to be a ubiquitously expressed key enzyme in cellular metabolism. This raises the question of how MMUT is regulated at the protein level, which may account for our observation of the two populations, MMUT+ and MMUT-. It has been shown that MMA leads to aberrant post-translational modifications by methymalonylation^60^. Moreover, novel phosphorylation sites in MMUT have been identified in liver samples^61^. Future work could look to identify the phosphoproteome during mitochondrial insult to identify a regulatory framework that leads to MMUT mislocalisation. (3) In acute excitotoxicity, the primary line of investigation centres around the idea that excessive postsynaptic depolarisation leads to cellular death, which may cause lesions seen in MMA but also in diseases such as Parkinson’s. However, we have no indication to believe that dopaminergic signalling is the reason behind basal ganglia lesions in MMA. Hence, excessive glutamate in the synapse from pre-synaptic cortical neurons innervating the striatum is just as likely a cause of excitotoxic cellular death and possibly cortical atrophy.

Here, we describe a mechanism of mitochondrial disruption generating reduced AP firing in patient-derived cells affected by MMA. Underlying this AP exhaustion was a reduced sodium current density, rewired glutamine metabolism, and differential glutamate receptor transcript expression. Further manipulation of glutamine metabolism reverses some of the metabolic and transcriptomic observations. More work will be required to elucidate the exact mechanism that reduces excitability in patient neurons.

## Methods

### Ethics statement, patients and fibroblast lines

Approval to use patient fibroblasts was granted by the Cantonal Ethics Commission of Zurich (KEK-ZH-Nr. 2014-0211, amendment PB_2020-00053). Both patient-derived fibroblast lines carried the *MMUT* homozygous pathogenic variant c.655A>T (p.Asn219Tyr), confirmed by Sanger sequencing at the DNA level, and corresponding to the mut^0^ phenotype. In addition to fibroblasts from two affected individuals (Pt1, female; Pt2, male), fibroblasts from Ct1 (ATCC: CRL-2522, male) were used for iPSC generation. An additional iPSC line control (Ct2, male; NHLBli003-A, RRID: CVCL_1E78) was acquired from Rutgers University (RUCDR, USA) ^62^.

### Fibroblast and HEK293T culture, and maintenance

HEK293T (ATCC: CRL-3216) and fibroblasts were cultured at 37°C and 5% CO2 in a humidified incubator. Cells were cultured in Dulbecco’s modified eagle medium (DMEM, Gibco, Cat. 31966-047) supplemented with 10% foetal bovine serum (FBS, Gibco, Cat. 102070-106) and 1% Anti-Anti (100X, Antibiotic-Antimycotic, Gibco, Cat. 15240-062). Passaging was performed once cells reached 80% confluency, using 0.05% Trypsin (1X, Trypsin-EDTA, Gibco, Cat. 25300-054) and washed with Dulbecco’s Phosphate Buffered Saline (DPBS, Gibco, Cat. 14190-250). Cells were maintained frozen in liquid nitrogen in cryopreservation medium (90% FBS (as before) and 10% DMSO (Sigma, Cat. D2438-50ML)).

### iPSC culture, differentiation, and maintenance

All cells were cultured at 37°C and 5% CO_2_ in a humidified incubator. The generation of iPSCs from fibroblasts was performed as previously described^63^. Briefly, the transduction was performed using the CytoTune®-iPS 2.0 Sendai Reprogramming Kit (Invitrogen, Cat. A16518) according to the manufacturer’s instructions. Transduction was performed on fibroblasts with passage numbers lower than five. Cells were plated at a density of 1×10^5^ cells/well in a 12-well plate. Non-integrating Sendai virus was added to the fibroblast medium at an MOI (ratio virus to cells) of 5 for KOS (Klf4, Oct4, Sox2) and c-Myc, and MOI 3 for Klf4. 8000 transduced fibroblasts were plated on inactivated mouse embryonic fibroblasts (MEFs). Derived iPSCs colonies were transferred to feeder-free conditions using 1mg/mL Collagenase type IV solution (Gibco, Cat. 17104019).

Feeder-free iPSC colonies were maintained as previously described^64^ with minor modifications. Between culturing, cells were maintained frozen in liquid nitrogen in cryopreservation medium (90% KnockOut™ Serum Replacement, Gibco, Cat. 10828028; 10% DMSO, Sigma, Cat. D2438). Cells were cultured on 0.17 mg/well (6-well) or 0.083 mg/well (12-well) BME matrix (Cultrex® Reduced Growth Factor Basement Membrane Extract, PathClear®, Cat. 3433-010-01) in Essential 8 medium (Gibco, Cat. A1517001) supplemented with 100 U/mL Penicillin-Streptomycin (10’000 U/mL, Gibco, Cat. 15140122). Passaging was performed once colonies reached 80% confluency using EZ-LiFT Stem Cell Passaging Reagent per manufactures instructions (Sigma-Aldrich, SCM139) after which 10 µL/mL RevitaCell™ Supplement (100X, Gibco, Cat. A2644501) was applied to passaged iPSC colonies. For neural induction, single cell passaging of iPSCs was performed using 0.5 mL or 1 mL ACCUTASE™ Cell detachment solution (1X, StemCellTechnologies, Cat. 07920) for 12-well and 6-well plates, respectively, and 2 µL/mL from 5 mM ROCK-inhibitor Y-27632 (Dihydrochloride; Sigma-Aldrich, Cat. SCM075) was applied for single cell passaging. Cells were inspected daily using the Leica DMIL microscope and bright-field culturing images were taken using the Leica Application Suite (LAS) Version 2.8.1 software.

### Array comparative genomic hybridization

To test the genetic integrity of fibroblast-derived iPSCs pre- and post-induction, DNA extracts from fibroblasts and iPSCs were assessed using array comparative genomic hybrid analysis (Cell Guidance Systems). Briefly, DNA was isolated from cell pellets using DNeasy Blood & Tissue Kit (Qiagen, Cat. 69504) per manufacturer’s instructions. All extracted DNA had a concentration >50 ng/µl and contained more than 1 µg. Purity was measured using an ND-1000 Spectrophotometer (NanoDrop Technologies, Inc.) and fell within A260/A230 = 1.8 – 2.2 and A260/A280 > 1.8. The array contained 754,027 distinct probes and analysis was conducted on the Infinium Global Screening Array v3.0 platform. The genome-wide resolution was ~200 kb. Copy number variations and loss of heterozygosity was reported in samples when present in at least ~15 – 20% of cells. The report can be found in Supplementary Figure 2.

### Germ-layer differentiation assay

iPSCs were differentiated to the three germ layers as previously described ^65,66^ with minor modifications. Cells were plated onto a 12-well plate in chemically defined medium with polyvinyl alcohol (CDM-PVA: 250 mL IMDM, LifeTechnologies Cat. 21980032); 250 mL DMEM/F-12 + GlutaMAX (Gibco, Cat. 31765027); 1 mL Concentrated Lipids (Life Technologies, Cat. 11905031); 20 µL 1-Thioglycerol (Sigma, Cat. M6145-25ML); 350 µL Insulin (Sigma, Cat. I9278-5ML); 250 µL Transferrin (Sigma, Cat. 10652202001); 10 mL PVA (10 g [stock: 0.05 g/mL], (Sigma, Cat. P8136-250G) in 200 mL embryo-grade water (Sigma, Cat. W1503)). For one day CDM-PVA was supplemented with 10 ng/mL Activin (Gibco, PHC9564) and 12 ng/mL fibroblast growth factor-2 (FGF2) (Gibco, Cat. PHG0024). Thereafter, lineage specific protocols were followed, as reported below.

For endoderm, 100 ng/mL Activin, 80 ng/mL FGF2, 10 ng/mL BMP-4 (Gibco, PHC9534), 10 μM LY294002 (Gibco, Cat. PHZ1144), and 3 μM CHIR99021 (Sigma, Cat. SML1046-5MG) were supplemented in CDM-PVA medium for one day. The next day, the medium was refreshed at the same concentrations without CHIR99021. On the third day, the medium was changed to RPMI medium (500 mL RPMI1640 + GlutaMAX (LifeTechnologies, Cat. 61870010); 10 mL B27 (Gibco, Cat. 17504044); 5 mL NEAA (Gibco, Cat. 11140035); 5 mL Pen/Strep (Gibco, Cat. 15140122)) containing 100 ng/mL Activin and 80 ng/mL FGF2. On day four cells were fixed with 4% paraformaldehyde (Sigma, Cat. 158127) in DPBS and were tested for markers of endoderm as described below.

For mesoderm, 100 ng/mL Activin, 20 ng/mL FGF2, 10 ng/mL BMP4, 10 μM LY294002, and 3 μM CHIR99021 was supplemented to CDM-PVA for two days, with media change each day. On day three cells were fixed with 4% paraformaldehyde in DPBS and were tested for markers of mesoderm as described below.

For ectoderm, 10 nM SB431542 (Sigma, Cat. S4317-5MG), 12 ng/mL FGF2, and 150 ng/mL Noggin (Sigma, Cat. H6416-10UG) was supplemented to CDM-PVA medium for ten days, with media change each day. On day eleven cells were fixed with 4% paraformaldehyde in DPBS and were tested for markers of ectoderm as described below.

To assess differentiation capacity, samples were analysed using the 3-Germ Layer Immunocytochemistry Kit (LifeTechnologies, Cat. A25538) per manufacturer’s instructions. TUJ1, AFP and SMA were included in this kit but anti-FoxA2/hHNF-Abs (diluted 1:100, R&D Systems, Cat. AF2400), anti-Brachyury-Abs (diluted 1:100, R&D Systems, Cat. AF2085), and anti-Nestin-Abs (diluted 1:100, Abcam, Cat. ab22035) were also acquired.

The next day, samples were incubated with the corresponding secondary Abs, which were diluted 1:250 in 1% BSA in PBS. AFP, SMA, and TUJ1 were counter-stained with AlexaFluor488 goat anti-mouse IgG1 (Invitrogen, Cat. A25536), AlexaFluor555 goat anti-mouse IgG2a (Cat. A25533), and AlexaFluor647 donkey anti-rabbit (Invitrogen, Cat. A25537), respectively from the 3-Germ Layer Immunocytochemistry Kit. FoxA2/hHNF and Brachyury were counter-stained with mouse-anti-goat Texas Red (Santa Cruz Biotechnology, Cat. sc-3916), and Nestin counter-stained with AlexaFluor 488 goat-anti mouse (Invitrogen, Cat. A21121). After the secondary Ab incubation, the samples were mounted using ProLong™ Diamond Antifade Mountant with DAPI (Invitrogen, Cat. P36971). Images were taken with the CLSM Leica SP5 resonant APD at the Center for Microscopy and Image Analysis, University of Zurich.

### Mature neuronal culture

To generate neural progenitors, iPSCs were disassociated into single cells using ACCUTASE™ Cell detachment solution as described above, and seeded at high density (1×10^6^ cells per well of a 12-well plate) in SMAD inhibition medium as previously described^27^. Cell counting was performed using the Countess 3 Automated Cell Counter (ThermoFisher Scientific) with trypan blue (Gibco, Cat. 15250061) staining to identify dead cells, as per manufactures instructions. Neural induction of iPSC was performed using the STEMdiff™ SMADi Neural Induction Kit (StemCellTechnologies, Cat. 08582) and protocol per manufacturer was followed^28^. Briefly, iPSCs were cultured for 8-12 days on BME matrix (as before) and passaged using 1 mg/mL Collagenase type IV solution (Gibco, Cat. 17104019) upon the appearance of neuroepithelial cells onto BME matrix coated 6-well plates in STEMdiff™ SMADi Neural Induction medium. The next day, medium was switched to neural maintenance medium (NMM: 48.4 mL KnockOut DMEM/F12, Gibco, Cat. 12660-102; 48.5 mL Neurobasal, Gibco, Cat. 12348-017; 1 mL B-27 Supplement, Gibco, Cat. 17504-044; 500 µL N-2 Supplement, Gibco, Cat. 17502-048; 500 µL Non-essential amino acids, Gibco, Cat. 11140-050; 500 µL GlutaMAX, Gibco, Cat. 35050-061; 500 µL Penicillin-Streptomycin, Gibco, Cat. 15140-122; 90 µL 2-Mercaptoethanol, Gibco, Cat. 31350010; 25 µL Insulin solution, Sigma, Cat. I9278-5ML; as previously described^27^) and henceforth medium was refreshed daily. Upon formation of rosettes (as described in^27,67^), cultures were treated with 5 µL/mL Recombinant Human Basic Fibroblast Growth Factor (bFGF, 4 µg/mL, Gibco, Cat. PHG0024). After four consecutive days of bFGF treatment, cells were passaged with 1 mg/mL Collagenase type IV solution and transferred to BME matrix (Cultrex®) coated 6-plates. Once morphologically apparent neuronal cells (polarised with leading axon and trailing dendrite) could be seen migrating from the apical border of the rosette, cultures were passaged with ACCUTASE™ Cell detachment solution and plated onto 10 μg/mL laminin (Sigma-Aldrich, Cat. L2020) and poly-L-Lysine (Sigma-Aldrich, Cat. P6282-5MG). Cells were passaged between day 21 and 26, and either continued as neural cultures or frozen for storage. For freezing, cells were detached using ACCUTASE™ Cell detachment solution, transferred to neural freezing medium (90% NMM, 10% DMSO (Sigma, Cat. D2438), 20 ng/mL bFGF), and placed in a Mr. Frosty (Thermo Scientific, Cat. 5100-0001) at −80°C and stored long-term in liquid nitrogen, as described^27^.

Neural cultures were fed daily with NMM until day 45, after which 48-hour cycles were implemented whereby every second day two thirds of the medium was removed and 2 mL NMM plus maturation supplements GDNF and BDNF (both at 20 ng/mL, StemCellTechnologies, Cat. 78058 and 78005) were supplied to neuronal cultures. In this way, neuronal cultures were maintained up to a maximum of day 120.

### Mitochondrial fluorometric stains and dyes

Mitochondrial oxidative phosphorylation complex I was inhibited using 1 µM rotenone (Sigma, Cat. R8875-1G) in DMSO for 30 minutes added directly to the medium. Mitochondrial membrane potential was assessed using tetramethylrhodamine-methyl-ester perchlorate (Sigma, Cat. T668) reconstituted in DMSO and supplemented at 15 nM incubated for 60 minutes. Mitochondrial superoxide production was measured using MitoSOX™ Mitochondrial Superoxide Indicators (Invitrogen, Cat. M36008) reconstituted in DMSO was supplemented at 5 µM for 15 minutes.

### Electrophysiology

Cells were used for electrophysiology studies after at least 50 days *in vitro* (range: 50 – 113, mean: 73) and at least 20 days following addition of the maturation supplements GDNF and BDNF in 10 μg/mL laminin (Sigma-Aldrich, Cat. L2020) and transfer to poly-L-Lysine (Sigma-Aldrich, Cat. P6282-5MG) coated Nunc™ Cell Culture/Petri Dishes (Thermo Scientific, Cat. 150318). For treatment studies, 0.1 mM, 6LmM or 12 mM dimethyl 2-oxoglutarate (Sigma, Cat. 349631) (DM-2OG) was added for 24 hours before cell collection. For recording, iPSC-derived neurons were placed in a recording chamber and whole-cell patch-clamp recordings were performed using an EPC10 USB amplifier (HEKA). iPSC-derived neurons were bathed with extracellular solution containing (in mM) 140 NaCl, 2 MgCl_2_, 2 CaCl_2_, 10 HEPES, 3 KCl, 10 D-Glucose, pH 7.4 (all from Sigma). Pipette solution contained (in mM) 4 NaCl, 120 K-gluconate, 10 HEPES, 10 EGTA, 3 Mg-ATP, 0.5 CaCl_2_, 1 MgCl_2_ pH 7.2 (all from Sigma). Patch pipettes ~9 MOhm were pulled from borosilicate glass capillaries (Harvard Apparatus, Cat. 30-0038). Voltages were corrected for a liquid junction potential of +12 mV. Series resistance was not compensated.

We used current-clamp recordings to examine AP firing and membrane properties. Step current injection protocols (duration, 2 s; step, 5 pA; −20 to +60 pA) were applied from the neuron’s resting membrane potential. Current-clamp data were filtered at 2.7 kHz and digitized with 20–100 kHz. To study whole-cell currents, voltage steps (duration, 1 s) were applied from a holding potential of −80 mV to voltages between −80 and +60 mV. Peak inward and outward currents were quantified and normalized to whole-cell capacitance to obtain current densities. Spontaneous excitatory postsynaptic currents were measured for 120 s at a holding potential of −80 mV. Voltage-clamp data were filtered at 2.9 kHz and digitized with 50 kHz.

Where described, MitoTracker™ Deep Red FM (Invitrogen, Cat. M22426) was used at 0.1 µM in NMM and provided directly to cells for 15 minutes. Thereafter, medium was aspirated, and cells washed with extracellular solution. Tested doses of MitoTracker™ Deep Red FM ≥ 1 µM were not compatible with electrophysiological recording.

Electrophysiological data were analysed with custom-written routines in IgorPro software (WaveMetrics, version 6.37) for current-clamp and voltage-clamp step protocols. Spontaneous synaptic events were analysed using miniML^68^ in Python (version 3.9). Extracted data were integrated using R Studio (version 2022.12.0+353). Analysis scripts are available upon request. Electrophysiology variables and groupings made during analysis are summarised in Supplementary Tables 1 & 2.

### Transcriptomics

Bulk RNA-seq was performed by Novogene from RNA extracted from liquid-nitrogen snap frozen cell pellets (DIV range: 65 – 78, mean: 75) that were lysed using QIAshredder (Qiagen, 79656) with RNA extracted using the RNeasy® Mini Kit (Qiagen, Cat. 74106). Prior to sequencing, RNA quality was assessed using an ND-1000 Spectrophotometer (NanoDrop Technologies, Inc.) to ensure sample purity was between A260/280 = 1.8-2.2 and A260/230 ≥ 1.8. Peak illustration was conducted using an Agilent 5400 and the low marker was used to calibrate the size of peaks. RNA libraries were prepared using poly(A) capture, followed by reverse transcription into cDNA. Sequencing was performed in PE150 (paired-end 150) mode on an Illumina NovaSeq 6000 at Novogene. Between 960 to 8800 ng of total RNA was provided and ≥ 20 million read pairs per sample were acquired.

Raw sequencing reads were preprocessed using the SUSHI framework ^69,70^, which was developed by the Functional Genomics Center Zurich (FGCZ). Adapter sequences and low-quality reads were trimmed off using fastp v0.02 ^70^. Filtered reads were pseudo-aligned against the reference human genome assembly GRCh38.p13 and quantified using Kallisto v0.46.1 ^71^. Differential gene expression analysis was performed between different conditions using the R package edgeR v3.42 ^72^. 21057 features were identified, and 15530 features had counts above the threshold. *P* < 0.05 (reported as false discovery rate unless otherwise stated, correction is done via Benjamini-Hochberg method) and one-fold change were considered to be differentially expressed. R package clusterProfiler was used for functional enrichment analysis based on Go terms and KEGG pathways, which utilised Over-Representation Analysis ^73^.

### Pseudo sc-RNAseq

To determine the cell type abundance, we used a deconvolution approach on the bulk gene expression profiles of the iPSC-derived neurons collected as described in Transcriptomics method section. We used the machine learning method CIBERSORTx^74^ to infer cell-type proportion without physical cell isolation. With CIBERSORTx^74^, we created a signature matrix of cell type identified in iPSC-derived neurons using published single-cell RNA-sequencing data by Gutierrez-Franco *et al*^75^, wherein the same iPSC differentiation protocol was used. Using this fine single-cell annotation of iPSC-derived neurons, we could run this digital cytometry tool on bulk transcriptomic expression to estimate cell type abundances.

### Metabolomics – Sugar phosphates

Polar metabolites were extracted from neuronal cells at day 60 cultured in 6-well plates, either untreated or after 24 hours supplementation with 12 mM DM-2OG, according to the described protocol^76^. After extraction, the metabolites were concentrated and dried via an Alpha 3-4 LSCbasic (Martin Christ, 102394, 102395) in vacuo centrifuge with an external oil-pump overnight at ambient temperature and stored at −80°C until analysis.

The metabolite extracts were dissolved in 100 μL of ultrapure water upon analysis, and a volume of 8 μL per sample was injected into an Agilent 1290 ultra-high performance liquid chromatography (UHPLC) module connect to an Agilent 6490A tandem quadrupole (QqQ) mass spectrometer, as previously described^32^. Chromatography was performed using a 0.25 μL/min flow rate and a gradient from 0 to 100% mobile phase B over a 25-minute total run time. Mobile phase A (12 mM acetic acid, 10 mM tetrabutylammonium, 2 mM acetylacetone, 3% methanol in mQ) and B (12 mM acetic acid, 10 mM tetrabutylammonium, 2 mM acetylacetone, 3% methanol, 80% acetonitrile in mQ) were used for phosphate sugar separation. The gradient method was as follows (time: % B): 11 min: 0%; 14 min: 15%; 19 min; 40%; 20 min; 100%; 21.5 minutes 0%. The QqQ MS was operated in dynamic multiple-reaction monitoring (MRM) mode. The list of transitions used to detect targeted compounds is reported in Supplementary Table 3. Skyline Software (v20.2, MacCoss Lab Software) was used for peak integration and resulting peak areas were normalized on total peak area ^32^. Data processing and statistical data analysis were performed in PRISM GraphPad (v.5.03). Technical triplicates were measured for each sample. Shapiro-Wilk test was used to assess normality. Significance was evaluated for multiple comparison via Kruskal-Wallis test with significance set at p<0.05 (without post hoc correction).

The normalization of the MS data for the TBA-based analysis was performed using the total peak area method, by dividing the area of the peak of interest by the sum of the areas of all peaks included in the analysis (as normalization factor), to derive relative abundances. The resulting relative abundances of the target compounds are thus unitless number, meaning a pure quantity without a physical unit.

This normalization method is widely used in MS-based metabolomics^32,77–80^, and it is particularly useful when the protein concentration of samples as estimation of the initial biomass cannot be determined due to the metabolite extraction protocol employed, as in the case of our study.

A complete list of metabolites detected, and their transition information, is available in Supplementary Table 3.

### Metabolomics – TCA cycle metabolites and stable isotope tracing

Treated cells were supplemented with 12 mM DM-2OG for 24 hours. Four hours before cell collection, medium was changed to NMM without GlutaMAX™ Supplement (Gibco, Cat. 35050061) and with 4 mM [U-^13^C] glutamine (Sigma-Aldrich, Cat. 605166). At collection, medium was removed, coverslips quickly dipped into sterile double-distilled water at 37 °C and quenched in 80% methanol at −20 °C. Cells were scrapped in methanol and centrifuged at 15,000g for 15Lminutes at 4 °C. Supernatants were collected, snap-frozen in liquid nitrogen and stored at −80 °C before LC–MS analysis. Mass spectrometry preparation and processing was performed as previously described ^10^.

### Immunocytochemistry

To detect native protein markers, immunochemical assay was performed on cells grown on glass coverslips treated with poly-L-Lysine. Depending on cell state, coverslips were also treated with BME matrix (iPSCs, NECs, NSCs) or laminin (NPCs, Neurons). Briefly, cells were washed with DPBS (Gibco, Cat. 14190144), and fixed with 4% paraformaldehyde (Sigma-Aldrich, Cat. 158127) in DPBS for 15 minutes at ambient temperature. Cells were then permeabilised for 30 minutes in 0.1% triton (diluted in DPBS), PFA was quenched using 100 mM glycine solution for 15 minutes, and coverslips blocked in 1% BSA for 60 minutes. Both primary and secondary antibodies were diluted in 1% BSA. All solutions were prepared in DPBS. A full list of antibodies, their sources, dilution ratios, incubation times and manufacturers can be found in Supplementary Table 4.

### Imaging and microscopy

Immunofluorescence images, unless otherwise stated, were taken with a DMi8 S Inverted Microscope fitted with a DFC480 camera (Leica) and RGB filter boxes. Image acquisition was adjusted for each fluorophore-antibody combination. All cells, unless otherwise stated, were plated on glass coverslips (Epredia, Cat. 630-2124) treated with poly-L-Lysine (Sigma-Aldrich, Cat. P6282-5MG) and mounted on Adhesive Microscopic Slides 25 x 75mm Menzel (ThermoFisher, Cat. J1800AMNZ) with either ProLong™ Gold Antifade Mountant with DNA Stain DAPI (Invitrogen, Cat. P36935) or ProLong™ Gold Antifade Mountant without DAPI (Invitrogen, Cat. P10144).

Images were also acquired using a Leica Confocal SP8 Inverse Falcon with Power HyD R photon detector in counting mode, and fluorophores excited using a white light laser or photomultiplier tube with a 405 diode to excite fluorophores in the UV spectrum. Images were acquired using an HC PL APO 63x/1,40 OIL CS2 objective (Leica) with oil immersion liquid (“Type-F” immersion liquid, Leica). Acquisition parameters for high resolution images and deconvolution: Z-step at 130 nm (Image slices range 18-25), Image size at 42.91 µm. Image deconvolution was performed using Bitplane Imaris.

### Correlation colocalization image analysis

The analysis of colocalization of the MMUT and TOMM20 channels was performed using the Coloc 2 plugin in ImageJ. This plugin performed a spatial pixel intensity correlation, creating a scatterplot with a linear regression fit, which indicated the degree of colocalization between the two channels. Pearson’s correlation analysis and the calculated Pearson’s R coefficient were extracted and reported, (https://imagej.net/Coloc_2 and https://imagej.net/Colocalization_Analysis).

### Immunoblotting

Cells were collected with Accutase or scraping, washed with DPBS, and centrifuged to retrieve pellets. The supernatant was removed, and the remaining pellet was either used immediately for the lysate production or stored at −80°C for a later use. The cell pellets were resuspended in 100 μl of lysis buffer containing 150 mM NaCl (Sigma, Cat.#S7653), 50 mM Trizma-Base (pH 8, Sigma, Cat.#T1503), 1% NP-40 (IGEPAL CA-630, Sigma, Cat.#I8896), 10% sodium deoxycholate (Sigma, Cat.#D6750) and 1% Halt™ Protease & Phosphatase Inhibitor Cocktail (100X, Thermo Fischer Scientific, Cat.#78440) and centrifuged max speed at 4°C for 5 minutes. The supernatant was collected in a pre-cooled microfuge tube. The protein concentration of each sample was assessed by performing a Bradford assay (Quick Start Bradford 1X Dye Reagent, BioRad, Cat.#5000205) and measuring the absorbance at 595 nm. The samples were prepared by adding the protein extract to a mixture of lysis buffer and 4X Laemmli buffer (BioRad, Cat.#161-0747) with 5% β-Mercaptoethanol (Sigma, Cat.#M6250) to obtain a total protein concentration of 1 μg/μL. The samples were incubated at 96°C for 5 minutes, before 15 µg was loaded, alongside a molecular weight standard (PageRulerTM Plus Prestained Protein Ladder, Thermo Fischer Scientific, Cat.#26619) on Novex™ Tris-Glycine Mini Protein Gels, 4–20%, 1.0 mm, WedgeWell™ format (Invitrogen, XP04200BOX) or self-cast 12% running gel (in µL: 2500 H_2_O, 3000 Bis-Acrylamide (Bio-Rad, Cat.#1610158) 30:0.8, 1875 1.5M Tris-HCl (pH 8.8), 75 10% SDS, 50 10% APS, 5 TEMED). The 10x running running buffer comprised 1.92 M glycine (PanReac AppliChem, Cat.#A1067,1000), 250 mM Trizma-Base and 35 mM Sodium dodecyl sulfate (SDS, Sigma, Cat.#L3771), which were dissolved in double-distilled water. The gels were run at 140V until the sample front reached the end of the gel. Proteins were transferred onto a nitrocellulose (pore size 0.45 µm; Cytiva, Cat.#10600007) or polyvinylidene fluoride (pore size 0.22 µm, activated in MeOH; Bio-Rad, Cat.#1620177) blotting membrane using the Trans-Blot® TurboTM Transfer Pack in running buffer + 20% methanol with Bio-Rad standard program (30 mins). Ponceau staining (Sigma, Cat.#P7170-1L) was performed to visualise protein transfer. Ponceau solution was washed off with distilled water and the membrane was blocked in blocking buffer at ambient temperature (10% 10x TBS-tween (24 g Tris base, 88 g NaCl (Sigma, Cat.#71380-1kG), in 1L distilled water neutralised to pH 7.6 with HCl) 0.2% Tween (Sigma, Cat.#P1379-100ML), 5% milk powder (Millipore, Cat.#70166-500G)) for 1 hour. Antibody staining was performed with agitation for 2 hours in ambient conditions or overnight at 4°C. Membranes were washed in 1xTBS-tween in between each step. Stained protein signal was developed using Cytiva GE ECL Solution (Amersham, RPN2109) or Clarity Max ECL solution (Bio-Rad, 1705062) on the ChemiDoc™ Touch imaging system (Bio-Rad). A full list of antibodies, their sources, dilution ratios, incubation times and manufacturers can be found in Supplementary Table 4. Uncropped gels can be found in Supplementary Figure 12.

### Propionate Incorporation Assay

The assay to quantify propionate pathway catabolism of iPSCs was assessed according to a protocol described previously^81^ and included modifications as described^26^.

### Quantitative PCR

To test for incorporation of remnants of the Sendai virus, RNA was extracted from fibroblasts, Sendai virus-positive fibroblasts and iPSCs using either the QIAamp RNA Blood Mini Kit (QIAGEN) or Qiagen RNeasy micro kit (QIAGEN) following manufacturer’s instructions. The isolated RNA was converted into cDNA, utilizing PrimeScript™ 1st strand cDNA Synthesis Kit (Takara, Cat.#6210A). cDNA was amplified by PCR, using Taq Polymerase (New England Biolabs) using the primers listed in Supplementary table 5. The PCR products were stained with EZ-Vision® One DNA Dye as Loading Buffer, 6X (Amresco), analyzed by gel electrophoresis on a 2% agarose gel and visualized by an UV transilluminator (Alpha Imager HP).

To test for expression of SOX2, PAX6 and EOMES, RNA was extracted from iPSC, neuroepithelium and NSCs as described above. cDNA was synthesised from 500-3000 ng RNA using the PrimeScriptTM II 1^st^ Strand cDNA Synthesis Kit using a Professional TRIO Thermocycler (Analytik Jena) following manufacturer’s instructions. Real-time quantitative PCR (RT-qPCR) consisted of 10 μL of GoTaq® qPCR Master Mix (2X, Promega, Cat.#A600A), 5 μL diethylpyrocarbonate (DEPC BioChemica, PanReac AppliChem, Cat.#A0881,0020) treated water and 1 μL of a 10 mM solution containing forward and reverse primers for the genes of interest, to which 4 μL of 300-3000 ng cDNA were added. The RT-qPCR was performed using either a LightCycler® 480 II (Roche), a 7900HT Fast Real-Time PCR System (Applied Biosystems), or a QuantStudio™ 7 Pro Flex Real-Time PCR System (Applied Biosystems, 4485701). In all cases the following conditions were used: 96°C for 5 minutes, followed by 45 cycles composed of 10 seconds at 96°C and 1 minute at 60°C. Technical triplicates were performed for each sample. Data were collected using the LightCycler® 480 Software release 1.5.0 SP4 or exported from the QuantStudio as CSV. The relative gene expressions were normalized using the housekeeping gene β-actin and analysed using the 2-^ΔΔCt^ method ^82^. Primer sequences can be found in Supplementary Table 5.

### Statistics and Reproducibility

Where necessary statistical testing is noted in the respective figure legend. Descriptive values and cell sources have been included in all figure legends. All datapoints have been displayed and what they represent is clearly stated.

## Data availability

All main data supporting the findings of this study are available within the article, Supplementary Data, and Supplementary Figures. RNA sequencing data have been deposed in Gene Expression Omnibus under the access number GSE289294. Source data can be found in Supplementary Data. Unprocessed blots can be found in Supplementary Figure 12. All other data are available from the corresponding author (or other sources, as applicable) on reasonable request.

## Acknowledgements

The authors thank Adhideb Ghosh for his assistance with transcriptomic data and Caroline Frei for her assistance with development of R analysis script. MCSD was supported by the FZK Fellowship of the University Children’s Hospital Zurich. DSF is supported by the Swiss National Science Foundation [310030_192505] and the University Research Priority Program of the University of Zurich ITINERARE – Innovative Therapies in Rare Diseases. MRB is supported by the Swiss National Science Foundation [310030_175779 and 310030_212505] and the University Research Priority Program of the University of Zurich ITINERARE – Innovative Therapies in Rare Diseases.

## Author contributions

MCSD, DSF, and MRB contributed to the conception and design of the study. MCSD carried out the experimental work contributing to the majority of the paper. MCSD and DSF prepared the manuscript with input from the other authors. MS generated patient-derived iPSCs. GM performed western blots and ICC. MAG performed western blots. DPR performed electrophysiology and ID helped with analysis and wrote some of the analysis scripts. MP performed TCA cycle metabolomics. FC performed HexP metabolomics. BV and FvM facilitated bulk transcriptomics and data integration. SC conducted bulk RNAseq deconvolution. All authors contributed to the manuscript and approved the submitted version.

## Competing interests

All authors declare no competing interests.

**Correspondence and requests for materials** should be addressed to D. Sean Froese, or Matthias R. Baumgartner

## References

1. Rath, S. et al. MitoCarta3.0: an updated mitochondrial proteome now with sub-organelle localization and pathway annotations. Nucleic Acids Res. 49, D1541–D1547 (2021).

2. Schaefer, A., Lim, A. & Gorman, G. Epidemiology of Mitochondrial Disease. in Diagnosis and Management of Mitochondrial Disorders (eds. Mancuso, M. & Klopstock, T.) 63–79 (Springer International Publishing, Cham, 2019). doi:10.1007/978-3-030-05517-2_4.

3. Rahman, S. Mitochondrial disease in children. J. Intern. Med. 287, 609–633 (2020).

4. Rahman, J. & Rahman, S. Mitochondrial medicine in the omics era. Lancet Lond. Engl. 391, 2560–2574 (2018).

5. Wajner, M. Neurological manifestations of organic acidurias. Nat. Rev. Neurol. 15, 253– 271 (2019).

6. DiMauro, S. & Schon, E. A. Mitochondrial disorders in the nervous system. Annu. Rev. Neurosci. 31, 91–123 (2008).

7. Chinnery, P. F. Primary Mitochondrial Disorders Overview. in GeneReviews® (eds. Adam, M. P. et al.) (University of Washington, Seattle, Seattle (WA), 1993).

8. Forny, P. et al. Guidelines for the diagnosis and management of methylmalonic acidaemia and propionic acidaemia: First revision. J. Inherit. Metab. Dis. 44, 566–592 (2021).

9. Froese, D. S. & Gravel, R. A. Genetic disorders of vitamin B12 metabolism: eight complementation groups – eight genes. Expert Rev. Mol. Med. 12, e37 (2010).

10. Forny, P. et al. Integrated multi-omics reveals anaplerotic insufficiency in methylmalonyl-CoA mutase deficiency. (2023) 10.1038/s42255-022-00720-8.

11. Kölker, S. et al. no Methylmalonic acid, a biochemical hallmark of methylmalonic acidurias but no inhibitor of mitochondrial respiratory chain. J. Biol. Chem. 278, 47388– 47393 (2003).

12. Luciani, A. et al. Impaired mitophagy links mitochondrial disease to epithelial stress in methylmalonyl-CoA mutase deficiency. Nat. Commun. 11, 970 (2020).

13. de Keyzer, Y. et al. Multiple OXPHOS deficiency in the liver, kidney, heart, and skeletal muscle of patients with methylmalonic aciduria and propionic aciduria. Pediatr. Res. 66, 91–95 (2009).

14. Collado, M. S. et al. Biochemical and anaplerotic applications of in vitro models of propionic acidemia and methylmalonic acidemia using patient-derived primary hepatocytes. Mol. Genet. Metab. 130, 183–196 (2020).

15. Wongkittichote, P. et al. Tricarboxylic acid cycle enzyme activities in a mouse model of methylmalonic aciduria. Mol. Genet. Metab. 128, 444–451 (2019).

16. Chandler, R. J. et al. Mitochondrial dysfunction in mut methylmalonic acidemia. FASEB J. 23, 1252–1261 (2009).

17. Manoli, I. et al. Targeting proximal tubule mitochondrial dysfunction attenuates the renal disease of methylmalonic acidemia. Proc. Natl. Acad. Sci. 110, 13552–13557 (2013).

18. Ramon, C., Traversi, F., Bürer, C., Froese, D. S. & Stelling, J. No Cellular and computational models reveal environmental and metabolic interactions in MMUT-type No methylmalonic aciduria. J. Inherit. Metab. Dis. **n/a**,.

19. Kandel, E. R., Koester, J. D., Mack, S. H. & Siegelbaum, S. A. Synaptic Transmission. In Principles of Neural Science, 6e (McGraw Hill, New York, NY, 2021).

20. Zheng, X. et al. Metabolic reprogramming during neuronal differentiation from aerobic glycolysis to neuronal oxidative phosphorylation. eLife 5, e13374 (2016).

21. Yellen, G. Fueling thought: Management of glycolysis and oxidative phosphorylation in neuronal metabolism. J. Cell Biol. 217, 2235–2246 (2018).

22. Bouzier-Sore, A.-K. et al. Competition between glucose and lactate as oxidative energy substrates in both neurons and astrocytes: a comparative NMR study. Eur. J. Neurosci. 24, 1687–1694 (2006).

23. Hall, C. N., Klein-Flügge, M. C., Howarth, C. & Attwell, D. Oxidative Phosphorylation, Not Glycolysis, Powers Presynaptic and Postsynaptic Mechanisms Underlying Brain Information Processing. J. Neurosci. 32, 8940–8951 (2012).

24. Forny, P. et al. Molecular Genetic Characterization of 151 Mut-Type Methylmalonic Aciduria Patients and Identification of 41 Novel Mutations in MUT. Hum. Mutat. 37, 745– 754 (2016).

25. Acquaviva, C. et al. N219Y, a new frequent mutation among mut° forms of methylmalonic acidemia in Caucasian patients. Eur. J. Hum. Genet. 9, 577–582 (2001).

26. Froese, S. & Baumgartner, M. Lysosomal Vitamin B12 Trafficking. in Ion and Molecule Transport in Lysosomes (CRC Press, 2020).

27. Shi, Y., Kirwan, P. & Livesey, F. J. Directed differentiation of human pluripotent stem cells to cerebral cortex neurons and neural networks. Nat. Protoc. 7, 1836–1846 (2012).

28. Chambers, S. M. et al. Highly efficient neural conversion of human ES and iPS cells by dual inhibition of SMAD signaling. Nat. Biotechnol. 27, 275–280 (2009).

29. Wurm, C. A. et al. Nanoscale distribution of mitochondrial import receptor Tom20 is adjusted to cellular conditions and exhibits an inner-cellular gradient. Proc. Natl. Acad. Sci. 108, 13546–13551 (2011).

30. Bardy, C. et al. Predicting the functional states of human iPSC-derived neurons with single-cell RNA-seq and electrophysiology. Mol. Psychiatry 21, 1573–1588 (2016).

31. Inak, G. et al. Defective metabolic programming impairs early neuronal morphogenesis in neural cultures and an organoid model of Leigh syndrome. Nat. Commun. 12, 1929 (2021).

32. van Scherpenzeel, M. et al. Dynamic tracing of sugar metabolism reveals the mechanisms of action of synthetic sugar analogs. Glycobiology 32, 239–250 (2022).

33. Fan, J. et al. GlutamineLdriven oxidative phosphorylation is a major ATP source in transformed mammalian cells in both normoxia and hypoxia. Mol. Syst. Biol. 9, 712 (2013).

34. Wu, G., Haynes, T. E., Li, H., Yan, W. & Meininger, C. J. Glutamine metabolism to glucosamine is necessary for glutamine inhibition of endothelial nitric oxide synthesis. Biochem. J. 353, 245–252 (2001).

35. Hansen, K. B. et al. Structure, Function, and Pharmacology of Glutamate Receptor Ion Channels. Pharmacol. Rev. 73, 298–487 (2021).

36. Hayasaka, K. et al. Comparison of Cytosolic and Mitochondrial Enzyme Alterations in the Livers of Propionic or Methylmalonic Acidemia: A Reduction of Cytochrome Oxidase Activity. Tohoku J. Exp. Med. 137, 329–334 (1982).

37. Martinelli, D. et al. Neurologic outcome following liver transplantation for methylmalonic aciduria. J. Inherit. Metab. Dis. **n/a**,.

38. Kaplan, P., Ficicioglu, C., Mazur, A. T., Palmieri, M. J. & Berry, G. T. Liver transplantation is not curative for methylmalonic acidopathy caused by methylmalonyl-CoA mutase deficiency. Mol. Genet. Metab. 88, 322–326 (2006).

39. Molema, F. et al. Neurotoxicity including posterior reversible encephalopathy syndrome after initiation of calcineurin inhibitors in transplanted methylmalonic acidemia patients: Two case reports and review of the literature. JIMD Rep. 51, 89–104 (2020).

40. Chakrapani, A., Sivakumar, P., McKiernan, P. J. & Leonard, J. V. Metabolic stroke in methylmalonic acidemia five years after liver transplantation. J. Pediatr. 140, 261–263 (2002).

41. Kasahara, M. et al. Current role of liver transplantation for methylmalonic acidemia: A review of the literature. Pediatr. Transplant. 10, 943–947 (2006).

42. Nyhan, W. L., Gargus, J. J., Boyle, K., Selby, R. & Koch, R. Progressive neurologic disability in methylmalonic acidemia despite transplantation of the liver. Eur. J. Pediatr. 161, 377–379 (2002).

43. Mc Guire, P. J., et al. Combined liver–kidney transplant for the management of methylmalonic aciduria: A case report and review of the literature. Mol. Genet. Metab. 93, 22–29 (2008).

44. Galera-Monge, T. et al. Mitochondrial Dysfunction and Calcium Dysregulation in Leigh Syndrome Induced Pluripotent Stem Cell Derived Neurons. Int. J. Mol. Sci. 21, 3191 (2020).

45. Jung, H.-Y., Mickus, T. & Spruston, N. Prolonged Sodium Channel Inactivation Contributes to Dendritic Action Potential Attenuation in Hippocampal Pyramidal Neurons. J. Neurosci. 17, 6639–6646 (1997).

46. Groten, C. J. & MacVicar, B. A. Mitochondrial Ca2+ uptake by the MCU facilitates pyramidal neuron excitability and metabolism during action potential firing. *Commun*. Biol. 5, 1–15 (2022).

47. Chang, D. T. W., Honick, A. S. & Reynolds, I. J. Mitochondrial trafficking to synapses in cultured primary cortical neurons. J. Neurosci. Off. J. Soc. Neurosci. 26, 7035–7045 (2006).

48. McKenna, M. C. The glutamate-glutamine cycle is not stoichiometric: Fates of glutamate in brain. J. Neurosci. Res. 85, 3347–3358 (2007).

49. Aldana, B. I. et al. Glutamate-glutamine homeostasis is perturbed in neurons and astrocytes derived from patient iPSC models of frontotemporal dementia. Mol. Brain 13, 125 (2020).

50. Andersen, J. V. et al. Deficient astrocyte metabolism impairs glutamine synthesis and neurotransmitter homeostasis in a mouse model of Alzheimer’s disease. Neurobiol. Dis. 148, 105198 (2021).

51. Shen, J. et al. Determination of the rate of the glutamate/glutamine cycle in the human brain by in vivo 13C NMR. Proc. Natl. Acad. Sci. 96, 8235–8240 (1999).

52. Chen, Q. et al. Rewiring of Glutamine Metabolism Is a Bioenergetic Adaptation of Human Cells with Mitochondrial DNA Mutations. Cell Metab. 27, 1007–1025.e5 (2018).

53. Yoo, H. C., Yu, Y. C., Sung, Y. & Han, J. M. Glutamine reliance in cell metabolism. Exp. Mol. Med. 52, 1496–1516 (2020).

54. Conte, F., van Buuringen, N., Voermans, N. C. & Lefeber, D. J. Galactose in human metabolism, glycosylation and congenital metabolic diseases: Time for a closer look. Biochim. Biophys. Acta BBA - Gen. Subj. 1865, 129898 (2021).

55. Lissin, D. V., Carroll, R. C., Nicoll, R. A., Malenka, R. C. & Zastrow, M. von. Rapid, Activation-Induced Redistribution of Ionotropic Glutamate Receptors in Cultured Hippocampal Neurons. J. Neurosci. 19, 1263–1272 (1999).

56. Lu, W.-Y. et al. Activation of Synaptic NMDA Receptors Induces Membrane Insertion of New AMPA Receptors and LTP in Cultured Hippocampal Neurons. Neuron 29, 243–254 (2001).

57. Mosbacher, J. et al. A Molecular Determinant for Submillisecond Desensitization in Glutamate Receptors. Science 266, 1059–1062 (1994).

58. Zhu, J. J., Esteban, J. A., Hayashi, Y. & Malinow, R. Postnatal synaptic potentiation: Delivery of GluR4-containing AMPA receptors by spontaneous activity. Nat. Neurosci. 3, 1098–1106 (2000).

59. Ballhausen, D., Mittaz, L., Boulat, O., Bonafé, L. & Braissant, O. Evidence for catabolic pathway of propionate metabolism in CNS: expression pattern of methylmalonyl-CoA mutase and propionyl-CoA carboxylase alpha-subunit in developing and adult rat brain. Neuroscience 164, 578–587 (2009).

60. Head, P. E. et al. Aberrant methylmalonylation underlies methylmalonic acidemia and is attenuated by an engineered sirtuin. Sci. Transl. Med. 14, eabn4772 (2022).

61. Bian, Y. et al. An enzyme assisted RP-RPLC approach for in-depth analysis of human liver phosphoproteome. J. Proteomics 96, 253–262 (2014).

62. Patterson, K., Linask, K. L., Beers, J. & Zou, J. Generation of two tdTomato reporter induced pluripotent stem cell lines (NHLBIi003-A-1 and NHLBIi003-A-2) by AAVS1 safe harbor gene-editing. Stem Cell Res. 42, 101673 (2020).

63. Chichagova, V., Sanchez-Vera, I., Armstrong, L., Steel, D. & Lako, M. Generation of Human Induced Pluripotent Stem Cells Using RNA-Based Sendai Virus System and Pluripotency Validation of the Resulting Cell Population. Methods Mol. Biol. Clifton NJ 1353, 285–307 (2016).

64. Ghaedi, M. & Niklason, L. E. Human Pluripotent Stem Cells (iPSC) Generation, Culture, and Differentiation to Lung Progenitor Cells. Methods Mol. Biol. Clifton NJ 1576, 55–92 (2019).

65. Vallier, L. & Pedersen, R. Differentiation of human embryonic stem cells in adherent and in chemically defined culture conditions. Curr. Protoc. Stem Cell Biol. **Chapter** 1, Unit 1D.4.1–1D.4.7 (2008).

66. Vallier, L. et al. Early Cell Fate Decisions of Human Embryonic Stem Cells and Mouse Epiblast Stem Cells Are Controlled by the Same Signalling Pathways. PLoS ONE 4, e6082 (2009).

67. Shi, Y., Kirwan, P., Smith, J., Robinson, H. P. C. & Livesey, F. J. No Human cerebral cortex development from pluripotent stem cells to functional excitatory synapses. Nat. Neurosci. 15, 10.1038/nn.3041 (2012).

68. O’Neill, P. S. et al. Deep learning-based synaptic event detection. 2023.11.02.565316 Preprint at 10.1101/2023.11.02.565316 (2023).

69. Hatakeyama, M. et al. SUSHI: an exquisite recipe for fully documented, reproducible and reusable NGS data analysis. BMC Bioinformatics 17, 228 (2016).

70. Chen, S., Zhou, Y., Chen, Y. & Gu, J. fastp: an ultra-fast all-in-one FASTQ preprocessor. Bioinforma. Oxf. Engl. 34, i884–i890 (2018).

71. Bray, N. L., Pimentel, H., Melsted, P. & Pachter, L. Near-optimal probabilistic RNA-seq quantification. Nat. Biotechnol. 34, 525–527 (2016).

72. Robinson, M. D., McCarthy, D. J. & Smyth, G. K. edgeR: a Bioconductor package for differential expression analysis of digital gene expression data. Bioinforma. Oxf. Engl. 26, 139–140 (2010).

73. Yaari, G., Bolen, C. R., Thakar, J. & Kleinstein, S. H. Quantitative set analysis for gene expression: a method to quantify gene set differential expression including gene-gene correlations. Nucleic Acids Res. 41, e170 (2013).

74. Newman, A. M. et al. Determining cell type abundance and expression from bulk tissues with digital cytometry. Nat. Biotechnol. 37, 773–782 (2019).

75. Gutiérrez-Franco, A. et al. Methanol fixation is the method of choice for droplet-based single-cell transcriptomics of neural cells. *Commun*. Biol. 6, 1–12 (2023).

76. Conte, F. et al. In Vitro Skeletal Muscle Model of PGM1 Deficiency Reveals Altered Energy Homeostasis. Int. J. Mol. Sci. 24, 8247 (2023).

77. Ruiz-Hernández, V., Roca, M. J., Egea-Cortines, M. & Weiss, J. A comparison of semi-quantitative methods suitable for establishing volatile profiles. Plant Methods 14, 67 (2018).

78. Sun, J. & Xia, Y. Pretreating and normalizing metabolomics data for statistical analysis. Genes Dis. 11, 100979 (2024).

79. Conte, F. et al. Isotopic Tracing of Nucleotide Sugar Metabolism in Human Pluripotent Stem Cells. Cells 12, 1765 (2023).

80. Guezenoc, J., Gallet-Budynek, A. & Bousquet, B. Critical review and advices on spectral-based normalization methods for LIBS quantitative analysis. Spectrochim. Acta Part B At. Spectrosc. 160, 105688 (2019).

81. Willard, H. F., Ambani, L. M., Hart, A. C., Mahoney, M. J. & Rosenberg, L. E. Rapid prenatal and postnatal detection of inborn errors of propionate, methylmalonate, and cobalamin metabolism: A sensitive assay using cultured cells. Hum. Genet. 34, 277–283 (1976).

82. Livak, K. J. & Schmittgen, T. D. Analysis of relative gene expression data using real-time quantitative PCR and the 2(-Delta Delta C(T)) Method. Methods San Diego Calif 25, 402–408 (2001).

