## Supplemental Material for "Mitochondrial dysfunction drives a neuronal exhaustion phenotype in methylmalonic aciduria"

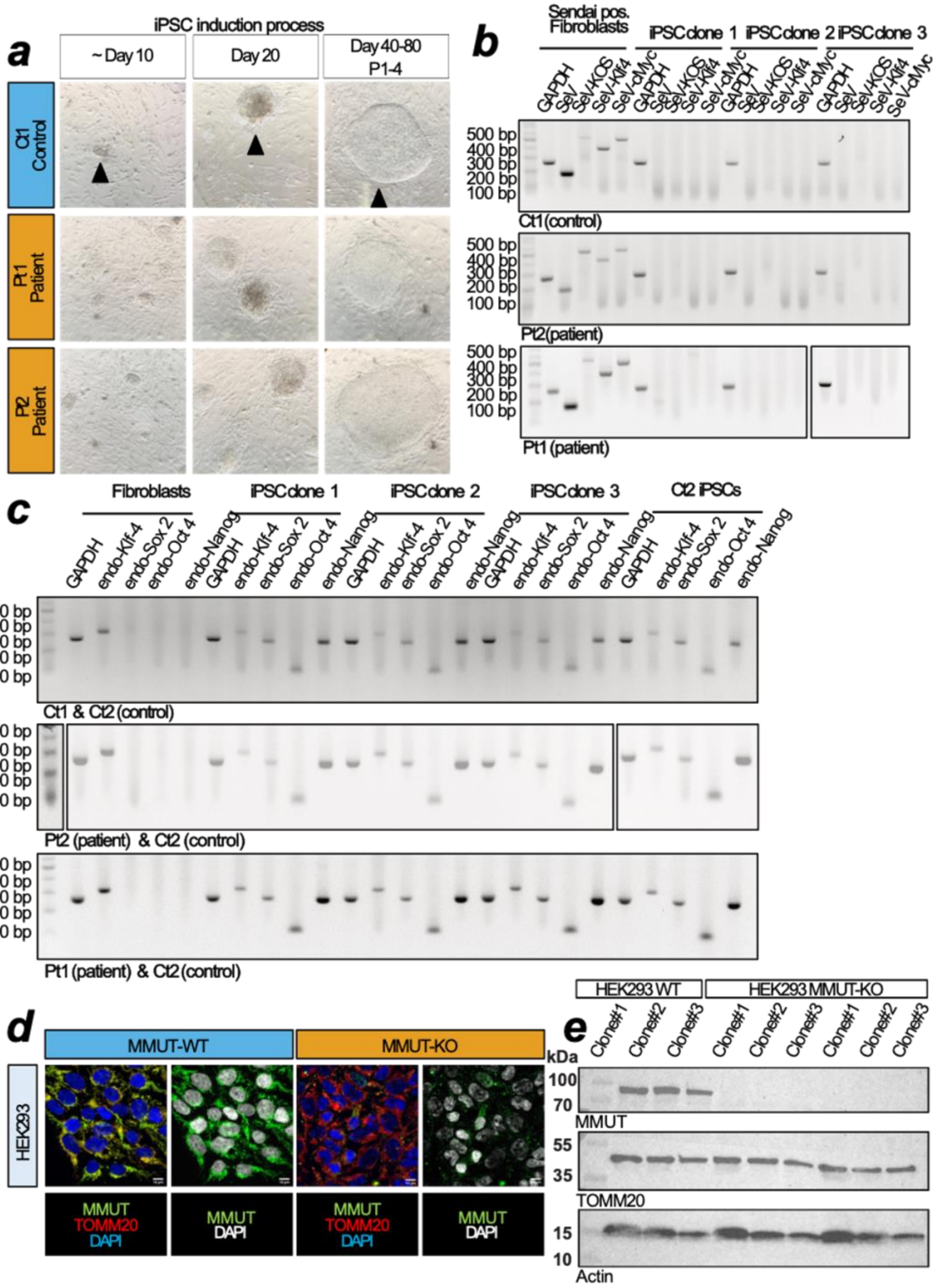


Supplementary Figure 1. **Fibroblasts generate iPSCs without incorporating viral factors. a.** Brightfield microscope images of iPSC generation process. **b.** Sendai virus clearance from fibroblasts to iPSCs in control (Ct1), and patient (Pt1, Pt2) cell lines. **c.** Endogenous mRNA expression of pluripotency markers in generated control, and patient cell lines, with purchased Ct2 control line. **d.** Immunochemistry of selected antibody against MMUT in WT and KO HEK293 cells. Scale bar = 10 µm. **e.** Western blot analysis of MMUT in WT and KO HEK293 cells. Uncropped membranes are available in Supplementary Data Fig. 2c.

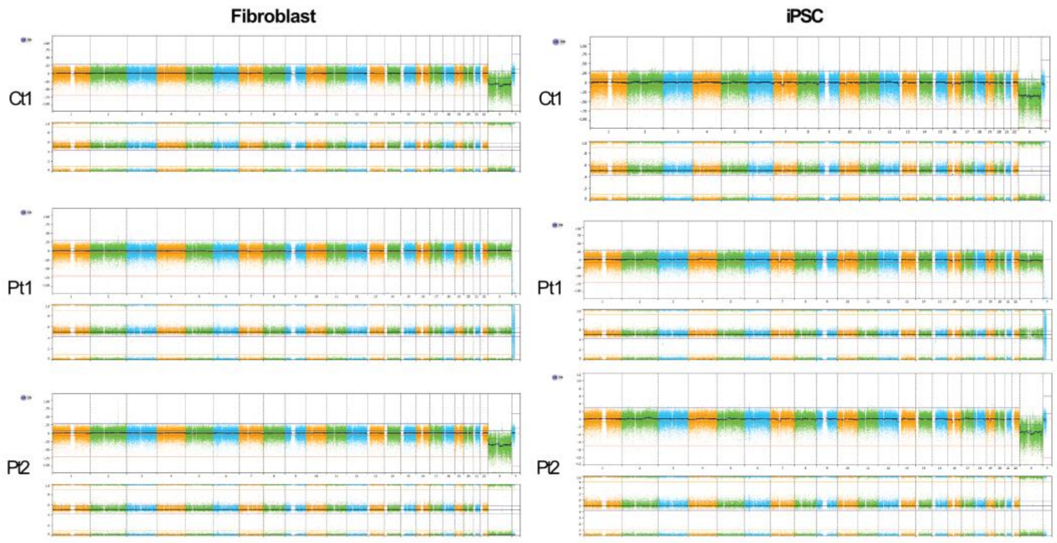


Supplementary Figure 2. Array comparative genomic hybridization (aCGH) analysis of fibroblasts and generated iPSCs. **a**, Visualization of aCGH to identify genomic rearrangements in one control (Ct1) and two patient (Pt1, Pt2) cell lines. Left are fibroblasts before pluripotency induction and right are after induction of pluripotency.

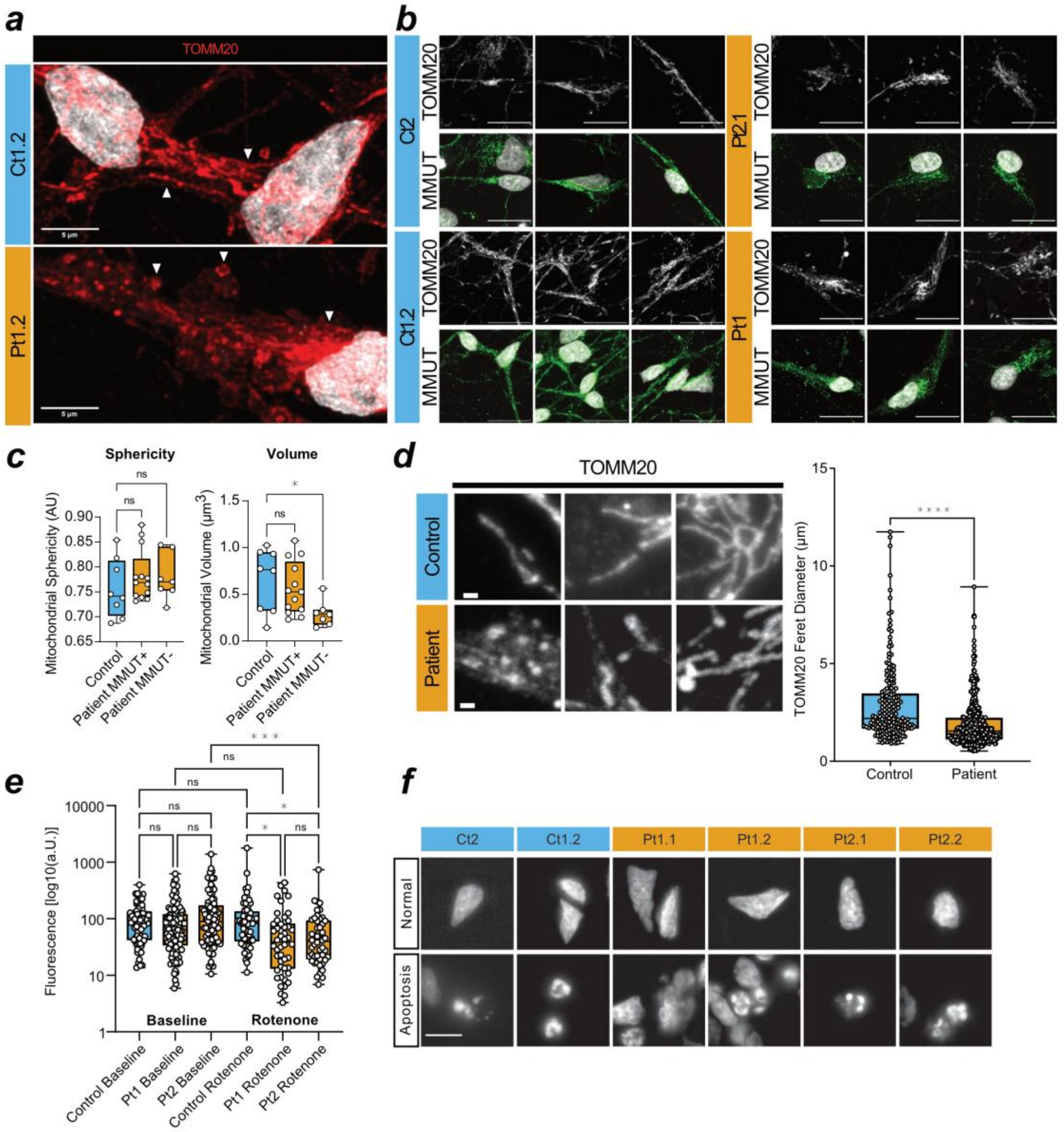


Supplementary Figure 3. **Patient-derived neurons have disrupted mitochondria. a.** Confocal microscope images from control and patient demonstrating TOMM20 staining pattern. Images are representative. Scale bar = 5 µm. **b.** Confocal images of localisation of MMUT (green) to TOMM20 (grey) in control (Ct1, Ct2) and patient (Pt1, Pt2) cell lines. Scale = 20 µm. **c.** Volume and sphericity measurements for Z-stack confocal images of TOMM20 in control and patient cells. Patient images were separated into MMUT localised (MMUT+) or MMUT mislocalised (MMUT-). Datapoints are representative of ROIs from one non-overlapping image. Datapoints represent Ct1 and Ct2 for control, and Pt1.1, Pt1.2, and Pt2.1 for patient. **d.** Maximum projection of confocal TOMM20 immunofluorescent images in control and patient (left), and ferret diameter measurements of those mitochondrial networks identified through TOMM20 staining. Scale =1 µm. Datapoints are representative of one ROI. Statistical test = Mann-Whitney, **** = <0.0001 (Right). Representative images and associated datapoints were selected from Ct2, Pt1.2, and Pt2.1. **e.** MitoSOX fluorescence measurements from confocal microscopy images in control and patient, without (left, baseline) and with (right, rotenone) 1 µM rotenone treatment. Datapoints are representative of one ROI. Statistical test = Kruskal-Wallis, ns = not significant, * =

<0.05, ** = <0.01,*** = <0.001. Datapoints represent Ct2 for control, and Pt1.2 and Pt2.1 for patient. **f.** Representative epifluorescence images of DAPI+ nuclei in control and patient during interphase and apoptosis. Scale = 10 µm.

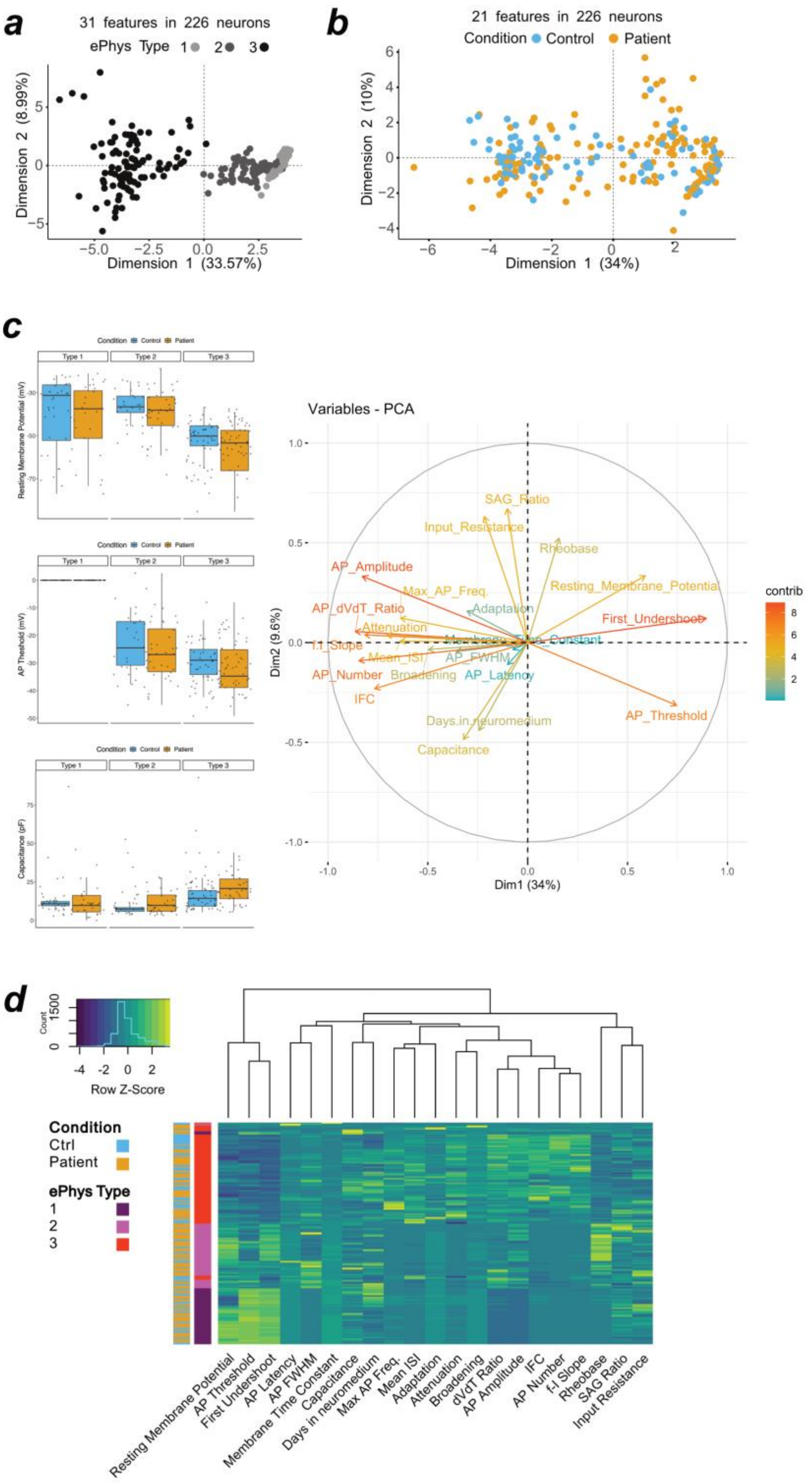


Supplementary Figure 4. **Electrophysiological characterisation of 226 neurons. a.** PCA of 31 features in 226 independent cells highlighting the 3 neuron types, measured during current-clamp recordings. List of features in Supplementary Data Table 1. **b.** Refined PCA of 21 features in 226 independent control (Ct1 n=59, Ct2 n=35) and patient (Pt1 n=77, Pt2 n=55) cells. **c.** Box plots indicate median and IQR 1.5 from Type 1-3 neurons for 3 selected features, each datapoint represents one independent cell (left). PCA loadings for the 21 features on 226 neurons in Fig. 4a, colour = percentage contribution to explained variance (right). **d.** Unguided hierarchical heatmap clustering on measured feature in 226 neurons. One row indicates one cell, one column represent one feature.

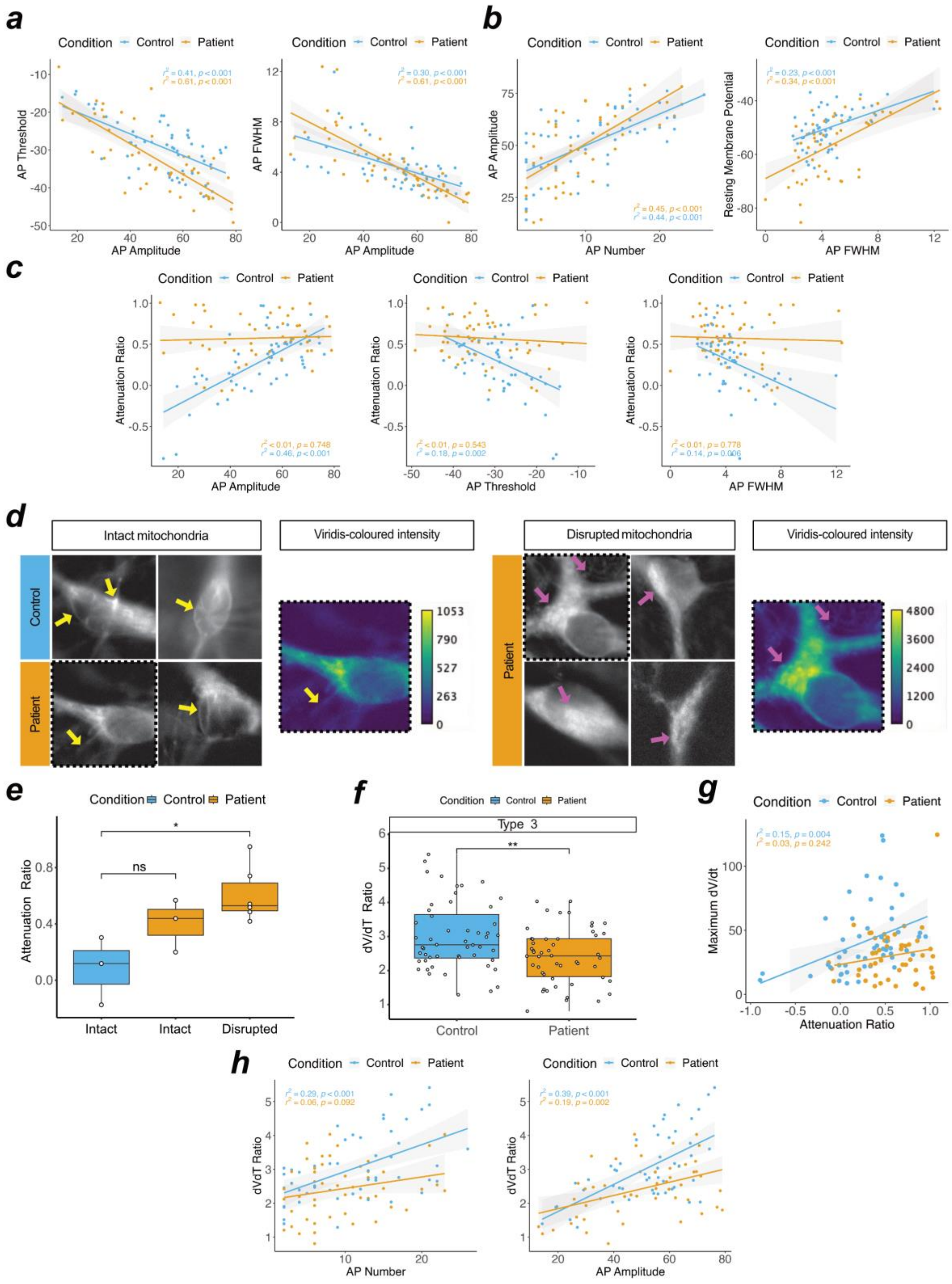


Supplementary Figure 5. **Loss of excitability in patient-derived neurons is explained by mitochondrial disruption. a.** Negative correlation between AP amplitude and threshold or full-width half-max in control and patient type 3 neurons. **b.** Positive correlation between AP amplitude and number or resting membrane potential and AP full-width half-max in control and patient type 3 neurons. **c.** Loss of correlation in patient-derived type 3 neurons between Attenuation ratio and AP amplitude, threshold or full-width half-max. **d.** Immunofluorescent images of MitoTracker in control and patient-derived neurons, showing intact (yellow arrowhead) or disrupted (magenta arrowhead) mitochondria, images are representative. Inset, pseudo coloured viridis-coloured images from representative cells demonstrate the relative fluorescent differences between intact and disrupted mitochondria. Units are arbitraty and histogram scale is match between images. Representative images were selected from Ct1, Pt1, and Pt2 **e.** Box plot indicates median and IQR 1.5 of attenuation ratio from 12 neurons measured in tandem with MitoTracker visualisation, each datapoint represents one independent cell (right). Datapoints represent data from Ct1, Pt1, and Pt2. **f.** Box plot indicates median and IQR 1.5 of dVdt in type 3 control and patient neurons, each datapoint represents one independent cell. **g.** Correlation between attenuation ratio and dVdt measurements in type 3 neurons, each datapoint represents one independent cell. **h.** Loss of correlation in patient-derived type 3 neurons between dVdt ratio and AP number or amplitude. Panel a-c, f-h feature 102 type 3 cells including control (Ct1 n=29, Ct2 n=23) and patient (Pt1 n=33, Pt2 n=17) cells.

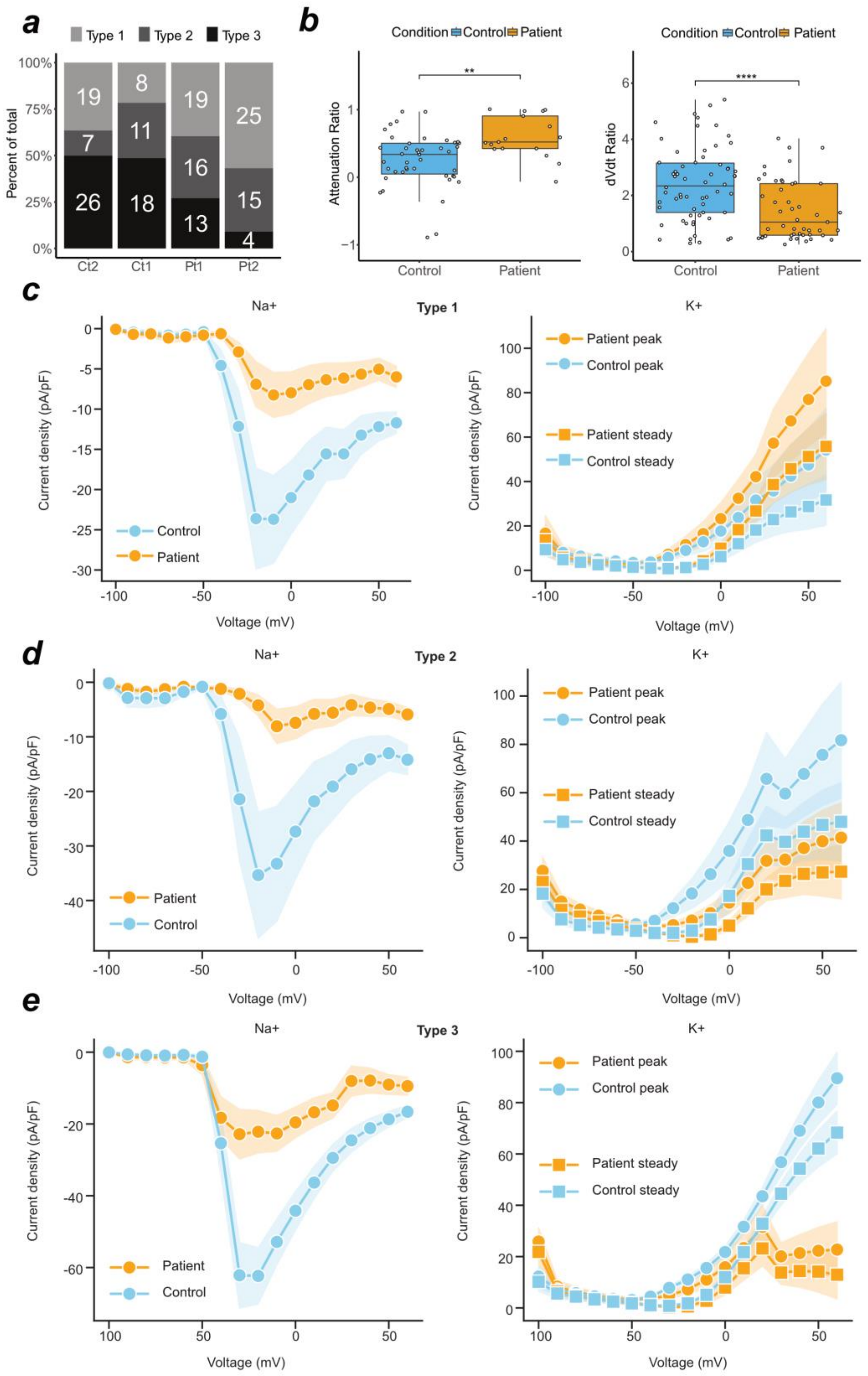


Supplementary Figure 6. **Attenuation phenotype in patient-derived neurons coherent with reduced sodium and potassium current densities a,** Stacked bar graph of proportional generation of type 1, 2, or 3 control (Ct1, Ct2) and patient (Pt1, Pt2) neurons in each cell line of cells measured in both current clamp and voltage clamp (n = 181). **b,** Box plot indicates median and IQR 1.5 of attenuation (left) and dVdt (right) ratios in control and patient neurons. **c,** Sodium and Potassium current density measurements from voltage-clamp recordings in 39 type 1 neurons. **d,** Sodium and Potassium current density measurements from voltage-clamp recordings in 42 type 2 neurons. **e,** Sodium and Potassium current density measurements from voltage-clamp recordings in 53 type 3 neurons.

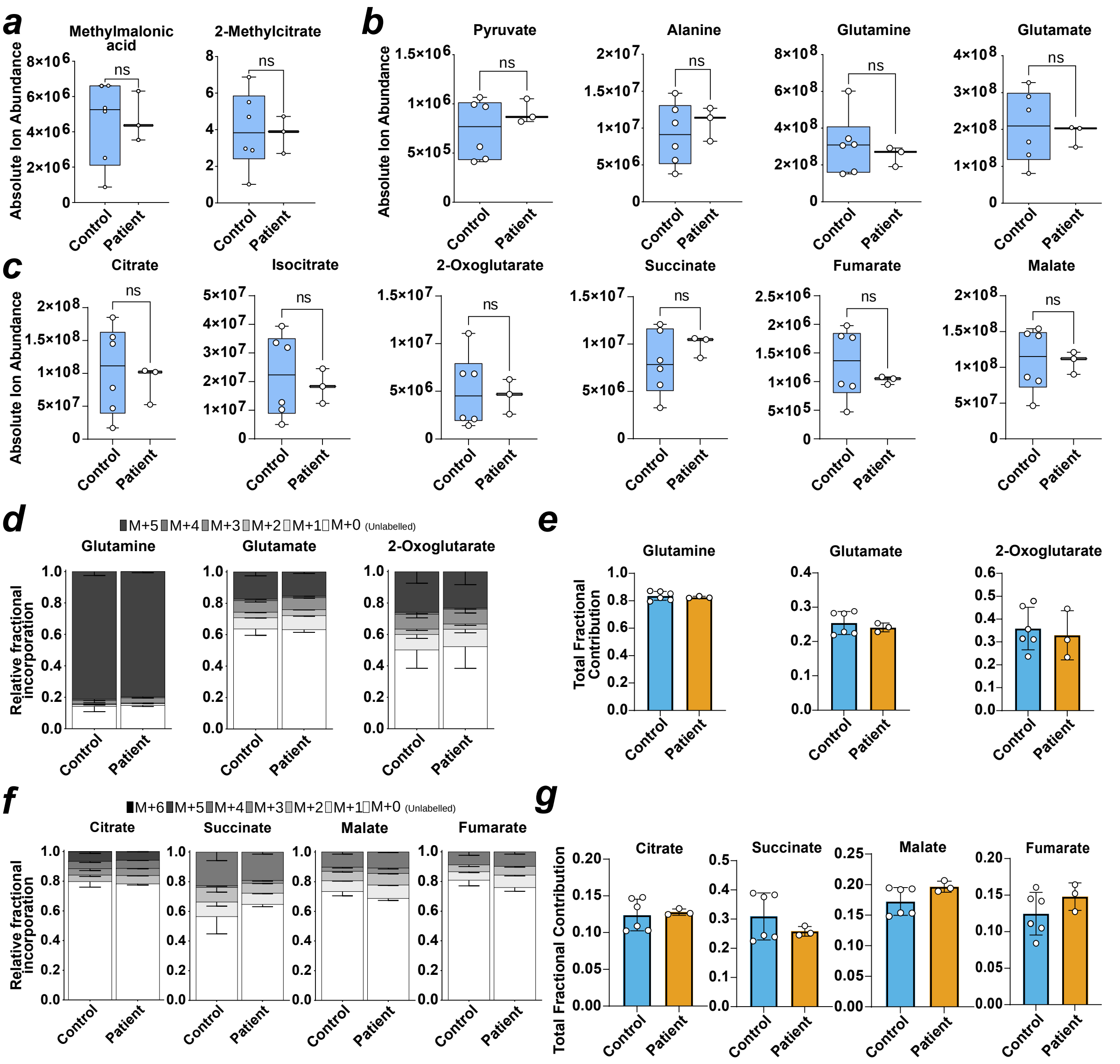


Supplementary Figure 7. **Absolute abundance of TCA cycle metabolites is unchanged in patient-derived neurons. a.** Box plot of ion abundance of disease-related metabolites. **b.** Box plot of ion abundance of anaplerotic metabolites. **c.** Box plot of ion abundance of TCA cycle metabolites. **d.** Relative fractional incorporation of fully labelled glutamine into glutamine, glutamate and 2-oxoglutarate. **e.** Bar plots of fractional contribution of labelled carbons to glutamine, glutamate and 2-oxoglutarate. **f.** Relative fractional incorporation of fully labelled glutamine into TCA cycle metabolites. **g.** Bar plots of fractional contribution of labelled carbons to TCA cycle metabolites. In **a, b, c, e, g,** each datapoint is an averaged result from technical triplicates of an independent sample. In **d, f** data is presented as a fraction of complete labelled and unlabelled measured metabolite. In **a-c**, significance = unpaired t-test. In all panels, datapoints represent Ct1.2 and Ct2 for control, and Pt1.2 and Pt2.2 for patient.

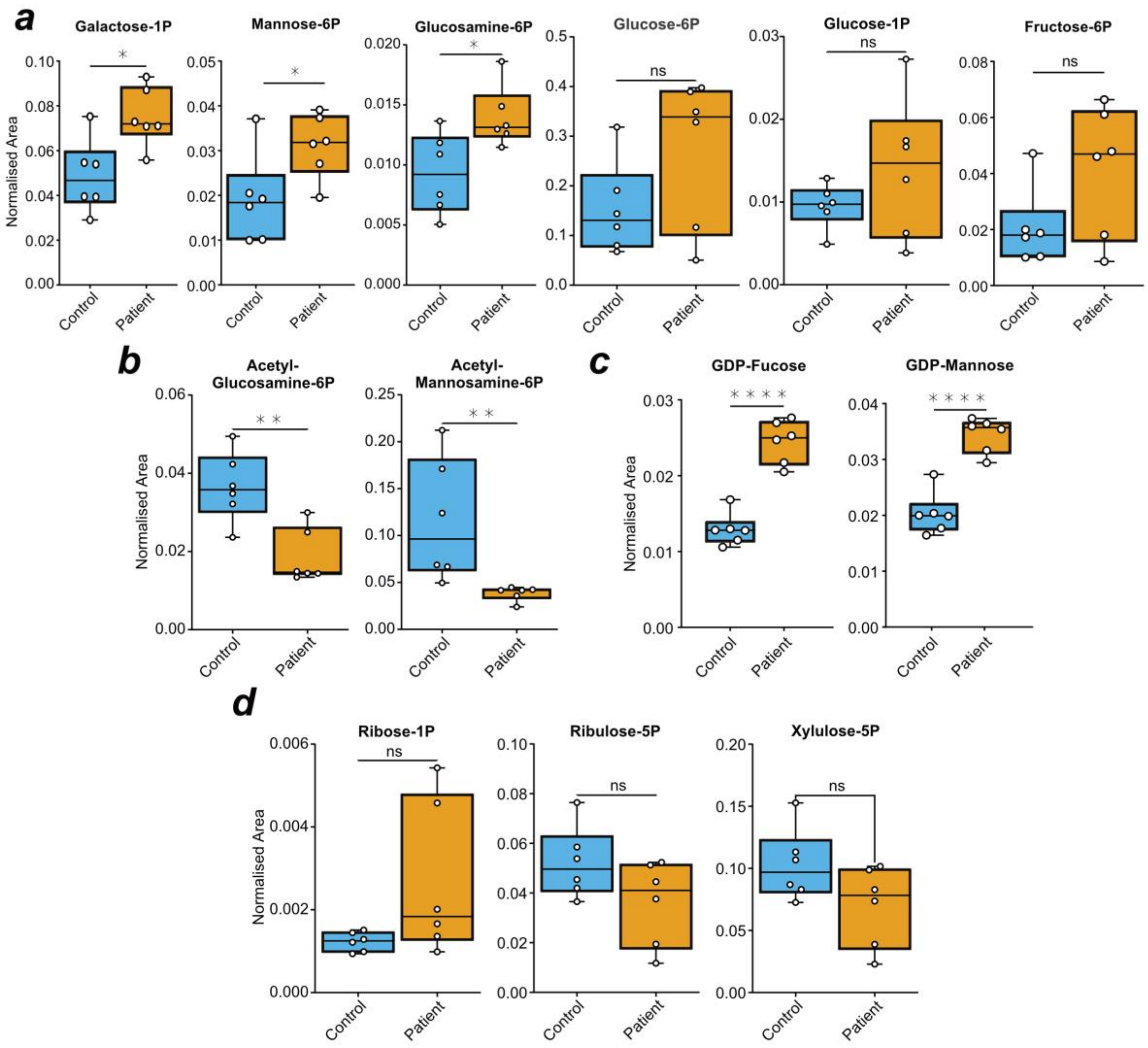


Supplementary Figure 8. **Changes in metabolite pools of selected sugar phosphates. a.** Box plot of normalised area of the peak of targeted hexose phosphates. **b.** Box plot of normalised area of the peak of amine-hexoses. **c.** Box plot of normalised area of the peak of targeted nucleotide sugars. **d.** Box plot of normalised area of the peak of targeted pentose phosphates. In **a-d,** significance testing = paired t-test. In all panels, datapoints represent Ct1.2 and Ct2 for control, and Pt1.2 and Pt2.2 for patient.

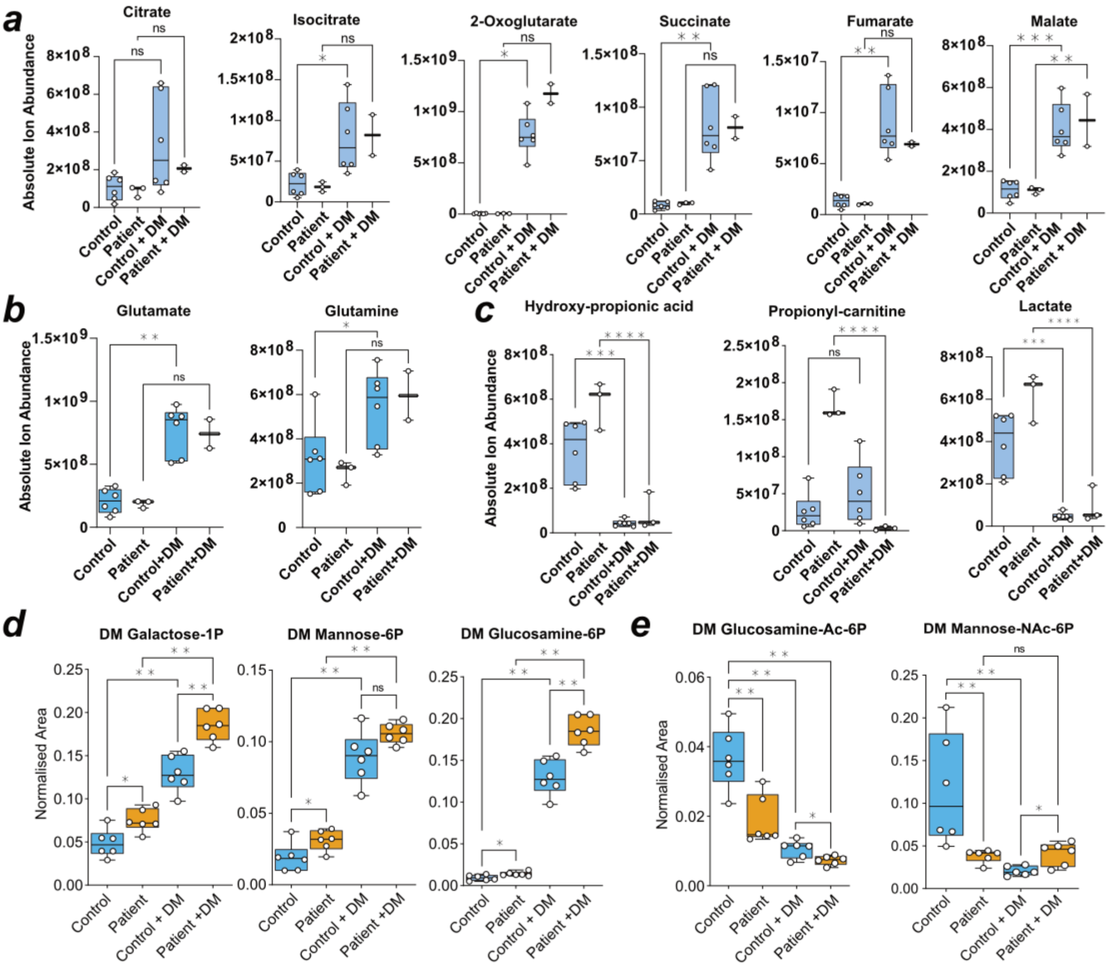


Supplementary Figure 9. **DM-2OG treatment rescues metabolic phenotype in patient neurons. a.** Box plot of response of absolute ion abundance of TCA cycle metabolites to treatment with dimethyl-2-oxoglutarate (DM). **b.** Box plot of glutamate and glutamine response to DM-2OG treatment. **c.** Box plot of response of absolute ion abundance of disease-related metabolites to treatment with dimethyl-2-oxoglutarate (DM). **d.** Box plot of normalised area of the peak of targeted hexoses and response to DM-2OG treatment. **e.** Box plot of normalised area of the peak of *N*-acetylhexosamine phosphates and response to DM-2OG treatment. In **a-e,** datapoints are representative of independent samples measured in triplicate and averaged. In **a-e** multiple testing = Mann-Whitney. In all panels, datapoints represent Ct1.2 and Ct2 for control, and Pt1.2 and Pt2.2 for patient.

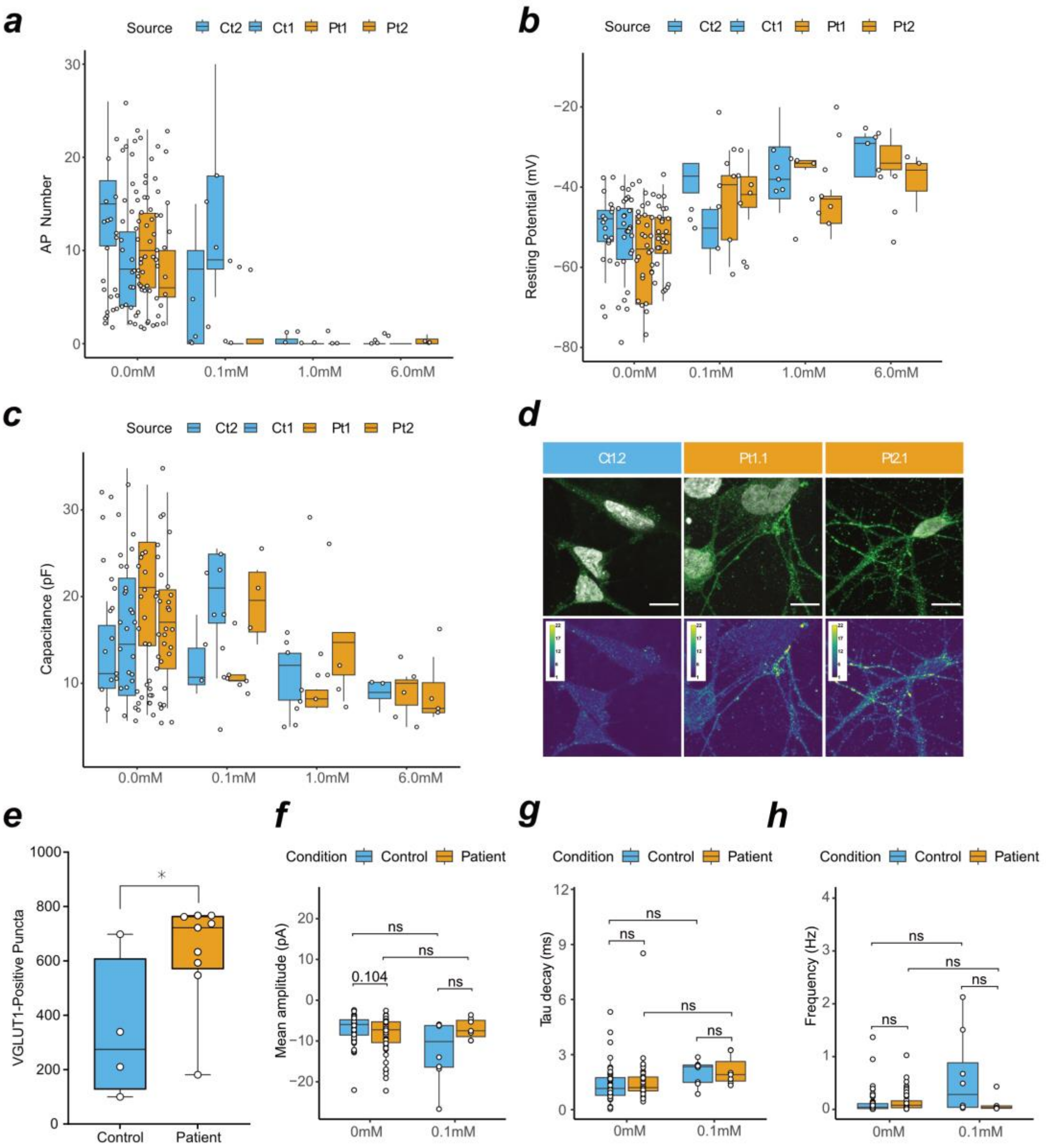


Supplementary Figure 10. **Patient excitatory perturbation can be mimicked in controls using DM-2OG. a.** Box plot of action potential count in current-clamp at various DM-2OG concentrations. **b.** Box plot indicates median and IQR at 1.5 from the measurement of resting membrane potential at different treatments of DM-2OG. **c.** Box plot indicates median and IQR at 1.5 from the measurement of capacitance at different treatments of DM-2OG. In **a, b, c,** datapoints are representative of an independently measured sample. **d.** Confocal immunofluorescence of an antibody against SLC17A7 (VGLUT1), images are representative. Scale = 10 µm. Representative images were selected from Ct1.2, Pt1.1, and Pt2.1. **e.** Box plot indicates median and min/max from quantification of synaptic VGLUT1 events in control (n=4) and patient (n = 9) neurons at DIV 50. Datapoints are representative of averages from non-overlapping images selected from Ct1.2, Pt1.1, and Pt2.1. **f.** Box plot of spontaneous event amplitude in response to DM-2OG treatment at 0.1mM. **g.** Box plot of spontaneous event decay in response to DM-2OG treatment at 0.1mM. **h.** Box plot of spontaneous event frequency in response to DM-2OG treatment at 0.1mM. In **f,g,h,** box plot indicates median and min/max. Where state numerically, $p$ value is Bonferroni significance adjusted. Datapoints are represent recordings from Ct1.2, Ct2, Pt1.1, Pt1.2, and Pt2.2.

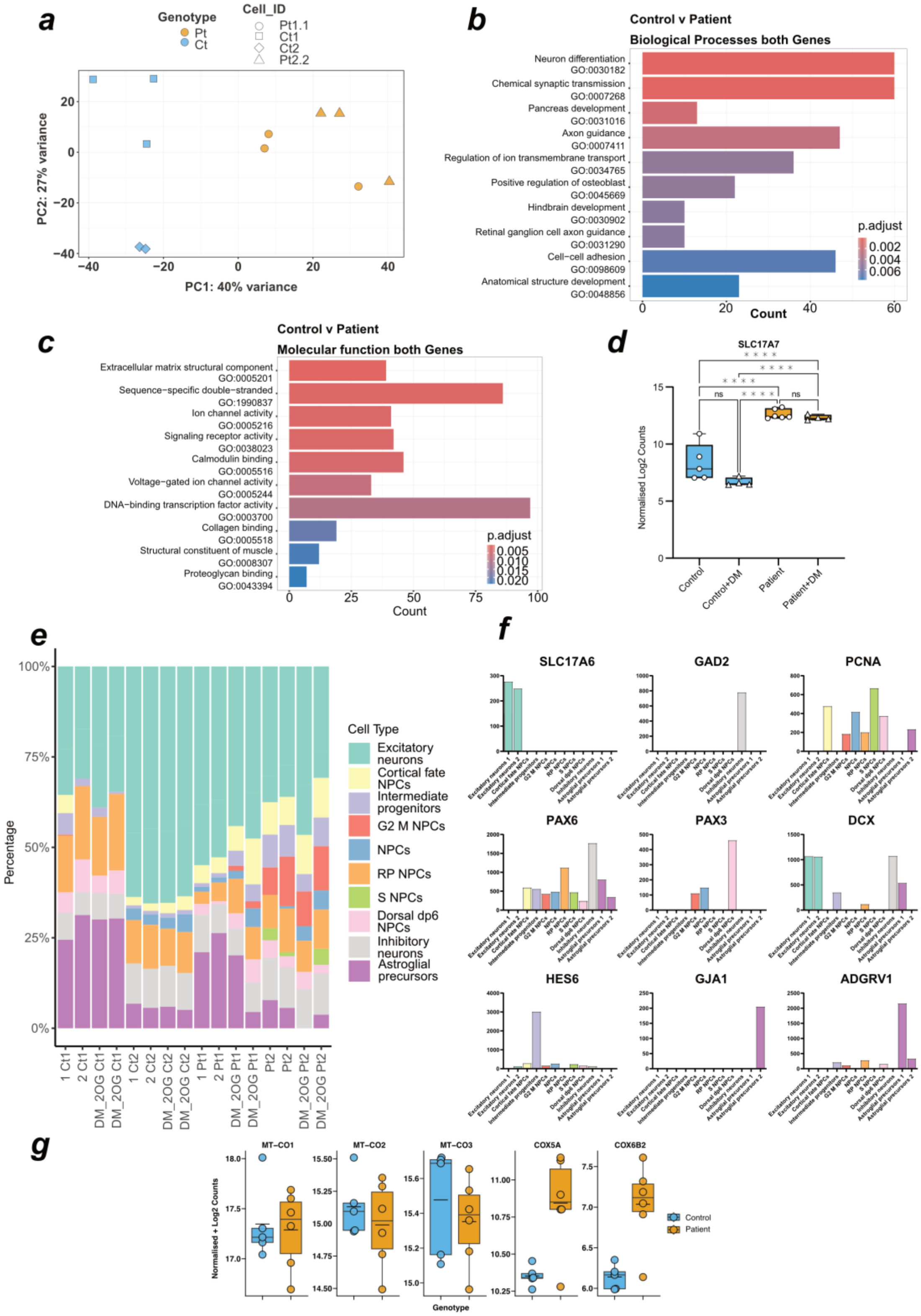


Supplementary Figure 11. **DM-2OG influences glutamate-sensitive synaptic apparatus. a.** PCA of control (n = 5) and patient (n = 6) neuronal cultures' transcriptome. **b.** Differentially regulated genes corresponding to GO terms associated to biological processes by overrepresentation analysis. **c.** Differentially regulated genes corresponding to GO terms associated to molecular functions by overrepresentation analysis. **d.** Box plot indicates median and min/max of count of the neuronal glutamate solute carrier extracted from transcriptomic data. Box plot indicates median and min/max of count of the selected genes extracted from transcriptomic data. **e.** Percentage of total transcripts mapped representing cell identity. Each column represents on independent cell sample. **f.** extracted transcripts for excitatory neurons (SLC17A6), inhibitory neurons (GAD2), proliferating marker (PCNA), dorsal fate markers (PAX6, PAX3), immature neuronal marker (DCX), intermediate progenitor markers (HES6), and astrocytic markers (GJA1, ADGRV1). **g.** Box plot indicates median and min/max, mean is overlaid bar of count of cytochrome c oxidase transcripts extracted from transcriptomic data. In **a, d,** datapoints are representative of an independently measured sample. In **b,c,** the top 10 (in either direction) differentially expressed terms were selected. In all panels, control is represented by Ct1.2 and Ct2, and patient is represented by Pt1.1, and Pt2.2.

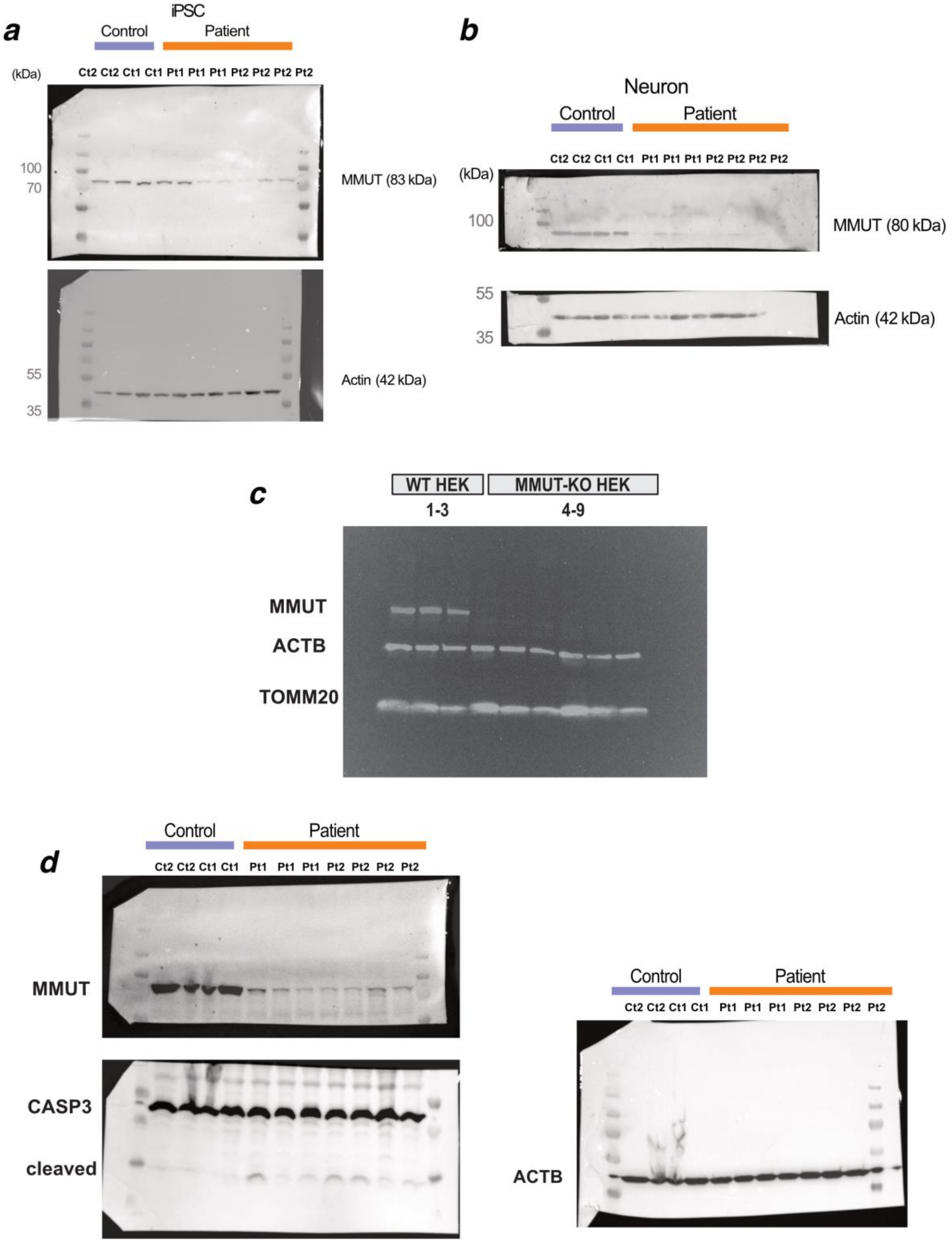


Supplementary Figure 12. Uncropped and unprocessed versions of western blots presented in the main and extended figures. **a,** MMUT protein in iPSCs. **b,** MMUT protein in differentiated neurons. **c,** MMUT protein in HEK cells. **d,** MMUT and CASP3 protein in differentiated neurons.

Supplementary Table 1. Electrophysiological variable collections and designations.

| Feature | Measurement type | Meaningful in ePhys type(s)... | Contrib. Ct2 | Contrib. Ct1 | Contrib. Pt1 | Contrib. Pt2 |
|---|---|---|---|---|---|---|
| **Rheobase** | Current_Clamp | T1-T3 | 35 | 59 | 77 | 55 |
| **First_Undershoot** | Current_Clamp | T2-T3 | 35 | 59 | 77 | 55 |
| **AP_Threshold** | Current_Clamp | T2-T3 | 35 | 59 | 77 | 55 |
| **Resting_Membrane_Potential** | Current_Clamp | T1-T3 | 35 | 59 | 77 | 55 |
| **AP_Latency** | Current_Clamp | T2-T3 | 35 | 59 | 77 | 55 |
| **Adaptation** | Current_Clamp | T3 | 35 | 59 | 77 | 55 |
| **Input_Resistance** | Current_Clamp | T1-T3 | 35 | 59 | 77 | 55 |
| **Mean_ISI** | Current_Clamp | T3 | 35 | 59 | 77 | 55 |
| **Membrane_Time_Constant** | Current_Clamp | T1-T3 | 35 | 59 | 77 | 55 |
| **SAG_Ratio** | Current_Clamp | T1-T3 | 35 | 59 | 77 | 55 |
| **AP_FWHM** | Current_Clamp | T2-T3 | 35 | 59 | 77 | 55 |
| **Max_AP_Freq.** | Current_Clamp | T2-T3 | 35 | 59 | 77 | 55 |
| **f.I_Slope** | Current_Clamp | T1-T3 | 35 | 59 | 77 | 55 |
| **AP_dVdT_Ratio** | Current_Clamp | T2-T3 | 35 | 59 | 77 | 55 |
| **dVdT_Maximum** | Current_Clamp | T2-T3 | 35 | 59 | 77 | 55 |
| **dVdT_Minimum** | Current_Clamp | T2-T3 | 35 | 59 | 77 | 55 |
| **IFC** | Current_Clamp | T3 | 35 | 59 | 77 | 55 |
| **Attenuation** | Current_Clamp | T3 | 35 | 59 | 77 | 55 |
| **Broadening** | Current_Clamp | T3 | 35 | 59 | 77 | 55 |
| **AP_Amplitude** | Current_Clamp | T2-T3 | 35 | 59 | 77 | 55 |
| **AP_Number** | Current_Clamp | T1-T3 | 35 | 59 | 77 | 55 |
| **Days.in.neuromedium** | Current_Clamp | T1-T3 | 35 | 59 | 77 | 55 |
| **Capacitance** | Current_Clamp | T1-T3 | 35 | 59 | 77 | 55 |
| **Batch** | Current_Clamp | T1-T3 | 35 | 59 | 77 | 55 |
| **Avg_AP_Freq.** | Current_Clamp | T2-T3 | 35 | 59 | 77 | 55 |
| **ISI_CV** | Current_Clamp | T3 | 35 | 59 | 77 | 55 |
| **Pause** | Current_Clamp | T3 | 35 | 59 | 77 | 55 |
| **Delay** | Current_Clamp | T3 | 35 | 59 | 77 | 55 |
| **Min_ISI** | Current_Clamp | T3 | 35 | 59 | 77 | 55 |

| | | | | | | |
|---|---|---|---|---|---|---|
| **Max_ISI** | Current_Clamp | T3 | 35 | 59 | 77 | 55 |
| **Median_ISI** | Current_Clamp | T3 | 35 | 59 | 77 | 55 |
| **Last_Undershoot** | Current_Clamp | T2-T3 | 35 | 59 | 77 | 55 |
| **Rectification** | Current_Clamp | T1-T3 | 35 | 59 | 77 | 55 |
| **Sodium Steady** | Voltage_Clamp | T1-T3 | 34 | 11 | 14 | 9 |
| **Sodium Peak** | Voltage_Clamp | T1-T3 | 34 | 11 | 14 | 9 |
| **Potassium Steady** | Voltage_Clamp | T1-T3 | 34 | 11 | 14 | 9 |
| **Potassium Peak** | Voltage_Clamp | T1-T3 | 34 | 11 | 14 | 9 |
| **Synaptic Event Frequency** | Voltage_Clamp | T1-T3 | 27 | 22 | 26 | 18 |
| **Synaptic Event Charge** | Voltage_Clamp | T1-T3 | 27 | 22 | 26 | 18 |
| **Synaptic Event Amplitude** | Voltage_Clamp | T1-T3 | 27 | 22 | 26 | 18 |
| **Mitotracker** | C_c & V_c | T1-T3 | 0 | 4 | 14 | 4 |
| **Dimethyl-2-Oxoglutarate** | C_c & V_c | T1-T3 | 17 | 5 | 13 | 12 |

Supplementary Table 2. Electrophysiological variable qualification into core groupings.

| | Variable | Core | Whole |
|---|---|---|---|
| Primary | **Rheobase** | **1** | **1** |
| | **First_Undershoot** | **1** | **1** |
| | **AP_Threshold** | **1** | **1** |
| | **Resting_Membrane_Potential** | **1** | **1** |
| | **AP_Latency** | **1** | **1** |
| | **Adaptation** | **1** | **1** |
| | **Input_Resistance** | **1** | **1** |
| | **Mean_ISI** | **1** | **1** |
| | **Membrane_Time_Constant** | **1** | **1** |
| | **SAG_Ratio** | **1** | **1** |
| | **AP_FWHM** | **1** | **1** |
| | **Max_AP_Freq.** | **1** | **1** |
| | **f.I_Slope** | **1** | **1** |
| | **AP_dVdT_Ratio** | **1** | **1** |
| | **IFC** | **1** | **1** |
| | **Attenuation** | **1** | **1** |
| | **Broadening** | **1** | **1** |
| | **AP_Amplitude** | **1** | **1** |
| | **AP_Number** | **1** | **1** |
| | **Days.in.neuromedium** | **1** | **1** |
| | **Capacitance** | **1** | **1** |
| Ancilliary | Batch | 0 | **1** |
| | Avg_AP_Freq. | 0 | **1** |
| | ISI_CV | 0 | **1** |
| | Pause | 0 | **1** |
| | Delay | 0 | **1** |
| | Min_ISI | 0 | **1** |
| | Max_ISI | 0 | **1** |
| | Median_ISI | 0 | **1** |
| | Last_Undershoot | 0 | **1** |
| | Rectification | 0 | **1** |

Supplementary Table 3. Metabolite transition list from acquisition method for hexose and pentose sugars.

| Compound Name | Precursor Ion | Product Ion | RT (min) | Delta RT | Fragmentor | Collision Energy | Cell Accelerator Voltage | Polarity |
|---|---|---|---|---|---|---|---|---|
| Glucosamine 6-phosphate | 258 | 79 | 1 | 3 | 380 | 45 | 7 | Negative |
| Glucosamine 6-phosphate | 258 | 97 | 1 | 3 | 380 | 15 | 7 | Negative |
| Glucosamine 6-phosphate | 258 | 227 | 1 | 3 | 380 | 7 | 7 | Negative |
| Glucosamine 6-phosphate | 258 | 240 | 1 | 3 | 380 | 9 | 7 | Negative |
| N-acetyl-D-glucosamine 6-phosphate / N-acetyl-D-mannosamine 6-phosphate | 300 | 79 | 12.8 | 3 | 380 | 48 | 7 | Negative |
| N-acetyl-D-glucosamine 6-phosphate / N-acetyl-D-mannosamine 6-phosphate | 300 | 97 | 12.8 | 3 | 380 | 15.5 | 7 | Negative |
| N-acetyl-D-glucosamine 6-phosphate / N-acetyl-D-mannosamine 6-phosphate | 300 | 282 | 12.8 | 3 | 380 | 6 | 7 | Negative |
| Fructose 6-phosphate - Galactose 1-phosphate - Glucose 1-phosphate - Glucose-6-phosphate - Mannose 6-phosphate | 259 | 79 | 10 | 7 | 380 | 50.5 | 7 | Negative |
| Fructose 6-phosphate - Galactose 1-phosphate - Glucose 1-phosphate - Glucose-6-phosphate - Mannose 6-phosphate | 259 | 97 | 10 | 7 | 380 | 13.3 | 7 | Negative |
| Fructose 6-phosphate - Galactose 1-phosphate - Glucose 1-phosphate - Glucose-6-phosphate - Mannose 6-phosphate | 259 | 139 | 10 | 7 | 380 | 10.8 | 7 | Negative |
| Fructose 6-phosphate - Glucose-6-phosphate - Mannose 6-phosphate | 259 | 169 | 10 | 7 | 380 | 5.8 | 7 | Negative |
| Galactose 6-phosphate - Glucose-6-phosphate - Mannose 6-phosphate | 259 | 199 | 10 | 7 | 380 | 7 | 7 | Negative |
| Ribose 1-phosphate / Ribulose 5-phosphate / Xylulose 5-phosphate | 229 | 79 | 12 | 6.5 | 380 | 40 | 7 | Negative |
| Ribose 1-phosphate / Ribulose 5-phosphate / Xylulose 5-phosphate | 229 | 97 | 12 | 6.5 | 380 | 9.2 | 7 | Negative |
| Ribose 1-phosphate / Ribulose 5-phosphate / Xylulose 5-phosphate | 229 | 139 | 12 | 6.5 | 380 | 10.25 | 7 | Negative |

Supplementary Table 4. Antibody list for immunochemistry.

| Target | Cat. No. | Manufacturer | Dilution<br>[ICC]<br>[WB] | Incubation<br>[ICC] (Temp.)<br>[WB] (Temp.) | Species |
|---|---|---|---|---|---|
| MMUT | Ab67869 | Abcam | 1:50<br>1:500 | Overnight 4°C<br>Overnight 4°C | Mouse |
| TOMM20 | Ab186735 | Abcam | 1:200<br>1:200 | 2 hours RT<br>Overnight 4°C | Rabbit |
| TOMM20 | sc11415 | Santa Cruz | 1:250<br>1:200 | 2 hours RT<br>Overnight 4°C | Rabbit |
| SOX2 | Ab5603 | Millipore | 1:400<br>- | Overnight 4°C<br>- | Mouse |
| SSEA4 | 53-8843-42 | Invitrogen | 1:200<br>- | 2 hours RT<br>- | Rabbit |
| NANOG | 4903S | Cell Signalling | 1:25<br>- | 2 hours RT<br>- | Rabbit |
| NANOG | 8750S | Cell Signalling | 1:50<br>- | 2 hours RT<br>- | Rabbit |
| Ki67 | Ab15580 | Abcam | 1:100<br>- | 2 hours RT<br>- | Rabbit |
| PAX6 | 901301 | Biolegend | 1:250<br>- | 2 hours RT<br>- | Rabbit |
| Nestin | Ab22035 | Abcam | 1:100<br>- | 2 hours RT<br>- | Mouse |
| TBR1 | Ab31940 | Abcam | 1:200<br>- | Overnight 4°C<br>- | Rabbit |
| Synaptophysin | Ab32127 | Abcam | 1:100<br>- | 2 hours RT<br>- | Rabbit |
| PSD-95 | Ab2723 | Abcam | 1:100<br>- | 2 hours RT<br>- | Mouse |
| NeuN | AB78 | Millipore | 1:500<br>- | Overnight 4°C<br>- | Rabbit |
| MAP2 | M1406-2ML | Sigma-Aldrich | 1:500<br>- | 2 hours RT<br>- | Mouse |
| EOMES | Ab23345 | Abcam | 1:100<br>- | 2 hours RT<br>- | Rabbit |
| TUBB3 | MAB1637 | Millipore | 1:200<br>- | 2 hours RT<br>- | Mouse |
| AFP | A25530 | Invitrogen | 1:100<br>- | Overnight 4°C<br>- | Mouse |
| FOXA2/hHNF | AF2400 | R&D | 1:100<br>- | Overnight 4°C<br>- | Goat |
| SMA | A25531 | Invitrogen | 1:50<br>- | Overnight 4°C<br>- | Rabbit |
| Brachyury | AF2085 | R&D | 1:100<br>- | Overnight 4°C<br>- | Goat |
| TUJ1 | A25532 | Invitrogen | 1:500<br>- | Overnight 4°C<br>- | Rabbit |
| TUBB3 | MAB1637 | MilliporeSigma | 1:250<br>- | 2 hours RT<br>- | Mouse |
| 20S ß5 | BML-PW8895-0025 | Enzo | 1:100<br>1:1000 | Overnight 4°C<br>Overnight 4°C | Rabbit |
| LC3 | 2775S | Cell Signaling Technologies | 1:100<br>1:1000 | Overnight 4°C<br>Overnight 4°C | Rabbit |
| Parkin | 66674 | ProteinTech | 1:100<br>1:1000 | Overnight 4°C<br>Overnight 4°C | Mouse |
| LAMP1 | H4A3-DSHB | Developmental Studies Hybridoma Bank | 1:100<br>1:1000 | Overnight 4°C<br>Overnight 4°C | Mouse |
| ß-Actin | A1978 | Sigma-Aldrich | -<br>1:1000 | -<br>Overnight 4°C | Mouse |
| SQSTM1 | 5114T | Cell Signalling Technologies | -<br>1:1000 | -<br>Overnight 4°C | Rabbit |
| Caspase-3 | 9662S | Cell Signalling Technologies | -<br>1:1000 | -<br>Overnight 4°C | Rabbit |
| VGLUT1 | 135303 | Synaptic Systems | 1:500<br>1:1000 | Overnight 4°C<br>Overnight 4°C | Rabbit |

| | | | | | |
|---|---|---|---|---|---|
| anti-mouse horseradish peroxidase | sc516102-cm | Santa Cruz | - 1:5000 | - 2 hours RT | Goat |
| anti-rabbit horseradish peroxidase | sc2357 | Santa Cruz | - 1:5000 | - 2 hours RT | Mouse |
| Alexa Fluor 594 F(ab') (H+L) | A11072 | Invitrogen | 1:1000 - | 1 hour RT - | Goat |
| Alexa Fluor 488 Cross-Adsorbed | A21121 | Invitrogen | 1:1000 - | 1 hour RT - | Goat |

Supplementary Table 5. Primer sequence list used in DNA PCR.

| Gene | FWD 5'-3' | REV 5'-3' | Product (bp) |
|---|---|---|---|
| **SOX2** | GAG CTT TGC AGG AAG TTT GC | TTC AAG GAG AGG CTT CTT GC | 190 |
| **PAX6** | AGC TCG GTG GTG TCT TTG TC | TGC ATC TGC ATG GGT CTG C | 128 |
| **EOMES** | GAG CCC TCA AAG ACC CAG AC | TCT GAA GCG GTG TAC ATG GAA | 155 |
| **ACTB** | AGG CAC CAG GGC GTG AT | GGC GTA CAG GGA TAG CAC AG | 318 |
| **GAPDH** | TGG ACC TGA CCT GCC GTC TA | ATG TGG GCC ATG AGG TCC ACC AC | 256 |
| **endo-Klf4** | ACC CAC ACA GGT GAG AAA CCT T | GTT GGG AAC TTG ACC ATG ATT G | 313 |
| **endo-Sox2** | TTC ATC GAC GAG GCT AAG CG | CAT CAT GCT GTA GCT GCC GT | 255 |
| **endo-Oct4** | GAG GAG TCC CAG GAC ATC AA | CAT CGG CCT GTG TAT ATC CC | 100 |
| **endo-Nanog** | CCA AAT TCT CCT GCC AGT GA | CAG GTG GTT TCC AAA CAA GAA | 260 |
| **SeV** | GGA TCA CTA GGT GAT ATC GAG C | ACC AGA CAA GAG TTT AAG AGA TAT GTA TC | 181 |
| **SeV-KOS** | ATG CAC CGC TAC GAC GTG AGC GC | ACC TTG ACA ATC CTG TAG TGG | 528 |
| **SeV-Klf4** | TTC CTG CAT GCC AGA GGA GCC | AAT GTA TCG AAG GTG CTC AA | 410 |
| **SeV-cMyc** | TAA CTG ACT AGC AGG CTT GTC G | TCC ACA TAC AGT CCT GGA TGA TGA TG | 532 |
